# A Two-Stage YOLO–Swin Transformer Pipeline for Cross-Dataset White Blood Cell Classification

**DOI:** 10.64898/2026.11.23.768502

**Authors:** Che-Wei Chang, Kuo-Jui Sun, Cheng-Hong Tsai, Chien-Yu Chen

## Abstract

**Background:** White blood cell (WBC) classification across datasets is challenged by differences in staining, image context, and class distributions.

**Methods:** We developed a two-stage YOLO11–Swin Transformer pipeline. In the first stage, WBC detection and context-preserving cropping, a YOLO11 detector finds the leukocyte in each uncropped image and produces a cell-focused crop that retains the surrounding background. In the second stage, WBC classification, a Swin Transformer (Swin-B) assigns the crop to one of nine cell types. The classifier was trained on detector-cropped cells from five public datasets, so the first stage supplies the same kind of input at test time; morphology-preserving color augmentation addressed staining variation, and class-weighted cross-entropy addressed class imbalance. WBCBench2026, a public benchmark not used for training, served for external evaluation in seven harmonized classes, scored by macro-F1 (the mean of per-class F1). Further analyses removed the first stage or masked the background, compared classifier backbones, held out each training dataset (leave-one-dataset-out, LODO), and tested a second split (Phase-2) with about 60% synthetically degraded images.

**Results:** The full pipeline reached an external macro-F1 of 0.879 ± 0.006 on 8,080 benchmark images. With uncropped images as input, the same crop-trained classifier reached 0.627 ± 0.039, and crops that kept the surrounding background outperformed background-masked crops (0.867 versus 0.760). With identical detector-cropped input, Swin-B outperformed convolutional baselines (0.863 versus 0.718–0.813), and its advantage over ResNet50 persisted across three training configurations. In LODO, macro-F1 on the held-out dataset ranged from 0.59 to 0.91, revealing source-specific variation not fully covered by the remaining datasets. On Phase-2, macro-F1 was 0.769 ± 0.002. Two hematologists independently reviewed all 205 blast- or promyelocyte-related model–benchmark disagreements and gave the same verdict on 73 crops (35.6%): 21 model right, 43 model wrong, 9 ambiguous.

**Conclusions:** The proposed two-stage pipeline achieved strong cross-dataset WBC classification performance. Context-preserving cropping was essential for the crop-trained classifier, and the LODO findings support representing the same classes across diverse sources when developing robust WBC classifiers.

## 1 Introduction

White blood cell (WBC) morphology assessment is central to hematology and to the diagnostic workup of leukemias, but manual differential counting is expert-dependent, labor-intensive, and shows appreciable inter-observer disagreement, both in the routine leukocyte differential^1^ and in the morphological criteria underpinning hematological diagnosis.^2^ Deep learning has therefore been proposed to standardize blood- and marrow-cell recognition, with human-level blast recognition reported^3^ and high precision and recall across marrow cell types,^4^ and now spans acute myeloid leukemia pathology from whole-slide-image mutation prediction^5^ to single-cell morphology analysis; automated WBC classification could support triage, education, and decision support, provided it generalizes across image sources.

Models developed on a single public WBC dataset frequently degrade when applied to images acquired under different staining protocols, microscopes or scanners, image resolutions, cropping conventions, and label taxonomies. This “dataset shift” is particularly consequential for minority or morphologically overlapping classes (e.g., promyelocytes and blasts), where a high overall accuracy can conceal poor class-specific performance. Such degradation across sites is well documented in radiology^6^ and in computational pathology, where scanner-and stain-related appearance shifts measurably reduce performance;^7, 8^ robustness under distribution shift is widely regarded as a prerequisite for clinical deployment.^9^

In the broader machine-learning literature this internal–external relationship has been characterized as “accuracy on the line”: across many models and natural distribution shifts, out-of-distribution accuracy correlates strongly and positively with in-distribution accuracy,^10–12^ so in-distribution accuracy is commonly treated as a proxy for deployed performance — although the correlation is weaker for some real shifts, such as tissue slides from different hospitals in the WILDS benchmark.^12, 13^

Most prior studies emphasize single-dataset accuracy; fewer evaluate external generalization, the role of the detection-and-cropping stage, or the sensitivity of a fixed-resolution classifier to inference-time changes. Whether transformer backbones are more robust than convolutional networks under external shift also remains incompletely characterized for cytomorphology: convolutional networks are the established WBC baseline, applied to normal peripheral-blood differentials^14^ and to blast recognition,^3^ whereas vision transformers have shown favorable robustness for image classification^15^ and have been benchmarked for medical imaging.^16^

This study develops a two-stage detect-then-classify pipeline — a trained YOLO11 detector^17, 18^ that passes a crop of its highest-confidence detection, with a 10% margin, to a Swin Transformer classifier^19^ — and makes the following contributions. (1) A cross-dataset WBC classification pipeline that integrates WBC detection and context-preserving cropping with multi-source classifier training on five public datasets and morphology-preserving color augmentation, evaluated as a complete system on a public external benchmark. (2) Ablations of the detection-and-cropping stage, augmentation and inference-time resolution: removing the detection-and-cropping stage collapsed external performance, a crop that kept the surrounding background outperformed one with the background masked, and the proposed color-augmentation recipe gave higher external macro-F1 and higher Blast group and Promyelocyte F1 than both comparator Swin-B recipes. (3) Back-bone comparisons in which Swin-B achieved higher external macro-F1 than ResNet50, by 0.058–0.169, under all three training configurations evaluated on both backbones, including a post hoc shared AdamW-based balanced parametric contrastive learning (BPaCo) protocol; internal macro-F1 on one common held-out set did not calibrate the level of external performance, and the two orderings inverted in some comparisons across training configurations. (4) A classifier-level leave-one-dataset-out (LODO) analysis showing that held-out sources contained variation not fully covered by the remaining sources, which supports representing the same cell classes across multiple data sources. (5) A coverage-aware evaluation of the complete pipeline, in which images the detector fails to localize are counted as errors rather than silently discarded, including on the partly degraded Phase-2 split, where a no-detection was a meaningful selective-prediction signal (Section 3.2). (6) A review of all 205 hardest-class model–benchmark disagreements (pre-cropped route; Section 3.9) by two hematologists who rated them independently: they gave the same verdict on 73 (21 model right, 43 model wrong and 9 ambiguous), indicating that expert-assigned benchmark labels can themselves contain errors.

## 2 Materials and Methods

The proposed pipeline has two stages (Fig. 1). The first, WBC detection and context-preserving cropping (below, the detection-and-cropping stage), runs a YOLO11 detector on the uncropped image and produces a cell-focused crop of the highest-confidence detection that retains its surrounding background; the second, WBC classification (the classification stage), classifies the crop with a Swin-B classifier trained on five public datasets with morphology-preserving color augmentation.

**Fig. 1.**
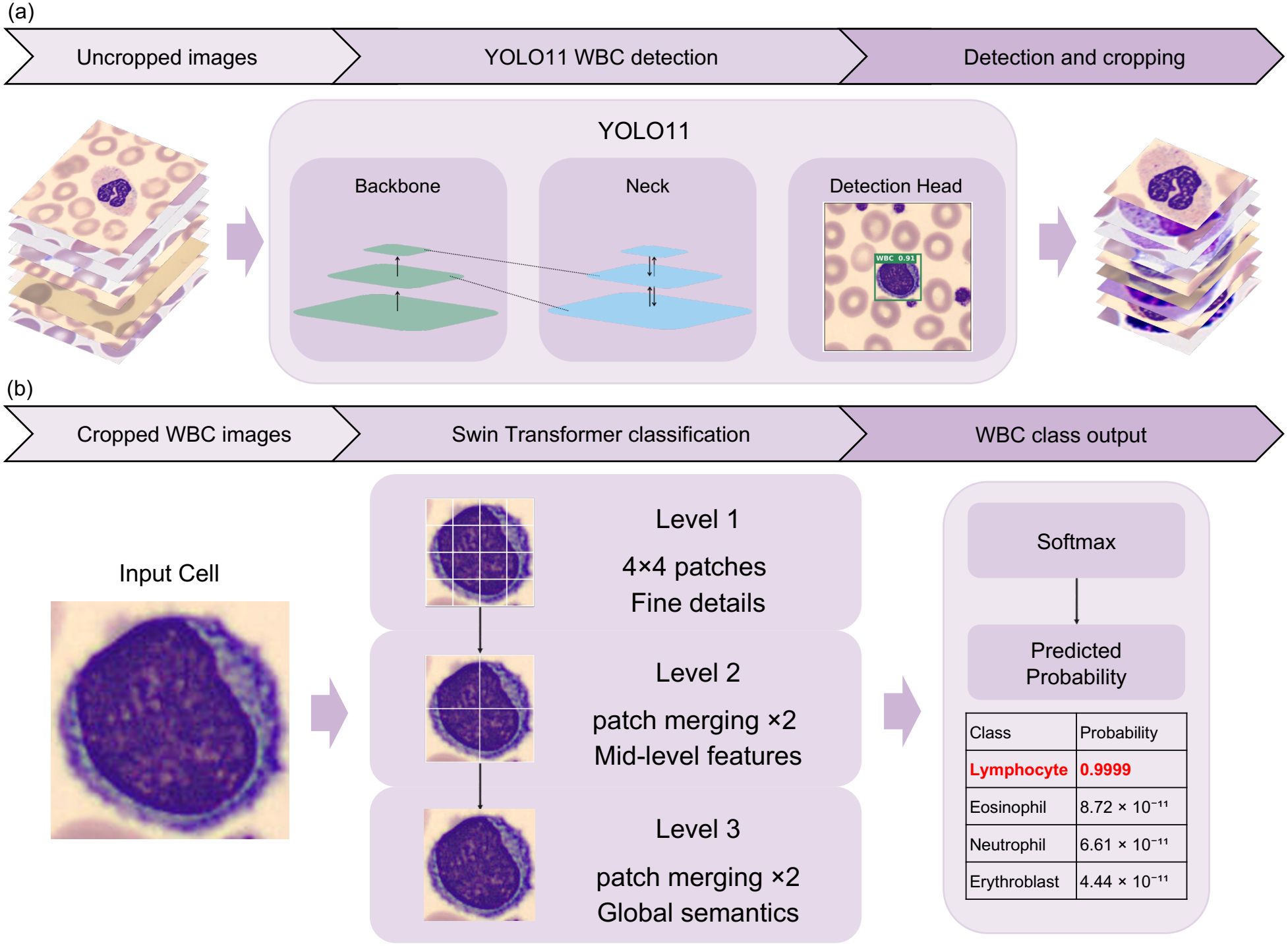
The two-stage YOLO–Swin white blood cell classification pipeline at inference. (a) Stage 1, WBC detection and context-preserving cropping: a trained YOLO11 detector finds the leukocyte in the uncropped image, and the highest-confidence detection, expanded by 10% on each side, is cropped with its surrounding background and resized to 224 × 224. (b) Stage 2, WBC classification: a Swin Transformer assigns the crop to one of nine cell types. The classifier was developed on five public datasets (127,938 single cells for training, validation and internal testing), and the WBCBench2026 Phase-1 split is used for external evaluation in a harmonized seven-class space. Example images are from PBC, AML-LMU, and HRLS; licenses are listed in Data availability.

### 2.1 Study design

This is a retrospective computational study of publicly available, de-identified WBC image datasets. For external evaluation we used the publicly released splits of the WBCBench2026 challenge dataset:^20, 21^ the Phase-1 release (8,288 images from 74 patients; each image, 368 px wide, contains one annotated leukocyte and is called an uncropped image below) served as the primary external evaluation split, and the released Phase-2 evaluation split (5,350 images from 49 patients not in Phase-1, about 60% of them with benchmark-defined synthetic acquisition degradation) served as a robustness test. Two routes through these images are reported. Classifier-level analyses (backbone and recipe comparisons, resolution, thresholds, calibration, error analysis and the hematologist review) use a pre-cropped set: the highest-confidence detector box per image at a threshold of 0.70 (as for the merged corpus; Section 2.2), with cells whose native labels have no counterpart in the output taxonomy removed, giving *n* = 8,061. The deployable full-pipeline route instead starts from all 8,288 uncropped images, runs the detector at 0.30, and counts an image with no detection as an error rather than dropping it; the same taxonomy exclusion removes 208 images, so *n* = 8,080 throughout (Table 2). The full attrition chain is in Supplementary Section S1, and we state the applicable *n* with every result. The challenge’s official test split was not used: its labels were not in the competition files used here. WBCBench2026 was not used for model training or checkpoint selection. Phase-1 performance informed the selection of augmentation and test-time augmentation (Supplementary Section S17); Phase-2 was evaluated after these choices were fixed.

### 2.2 Datasets

The classifier was trained and internally evaluated on five public WBC datasets (Table 1), and a further public detection dataset (TXL-PBC) was used to train the detector. The five training sources are referred to through-out by the short names defined in Table 1: AML-LMU,^22^ BMCMHF,^4, 23^ PBC^24^ (distinct from the TXL-PBC detection dataset below), Raabin-WBC,^25^ and HRLS.^26^ TXL-PBC^27^ integrates four public sources, two of which (PBC and Raabin-WBC) also appear among the classifier training datasets; WBCBench2026 is neither and postdates them, so by provenance its images were unseen by both pipeline stages during training, and a content-hash audit confirms that none of the detector-training images appears among them (Supplementary Section S1).

**Table 1.** Datasets used for classifier development, external evaluation and detector training. “Classes (source / used)”: categories in the released dataset / categories mapped into the label space used here (nine internal classes; seven evaluation classes for WBCBench2026; WBC only for TXL-PBC; Section 2.3). “Development”: classifier training, validation and internal testing. “Pool size”: each training dataset’s harmonized single-cell pool, used directly by the leave-one-dataset-out analysis; the five pools sum to 128,628, and the merged development corpus contains 127,938 cells (difference reconciled image by image in Supplementary Section S1). For WBCBench2026 the count is the released Phase-1 split and for TXL-PBC a field-image count. Specimen and staining are from each dataset’s primary publication. PB, peripheral blood; BM, bone marrow; MGG, May–Grünwald–Giemsa; MG, May–Grünwald.

| Dataset | Role | Pool size | Classes (source / used) | Specimen | Staining / notes |
|---|---|---|---|---|---|
| AML-LMU<sup>22</sup> | Development | 18,128 | 15 / 8 | PB | Not reported |
| BMCMHF<sup>4, 23</sup> | Development | 73,096 | 21 / 7 | BM | MGG (Pappenheim) |
| PBC<sup>24</sup> | Development | 11,845 | 8 / 6 | PB | MGG |
| Raabin-WBC<sup>25</sup> | Development | 10,154 | 5 / 5 | PB | Giemsa |
| HRLS<sup>26</sup> | Development | 15,405 | 9 / 7 | PB | MG + Giemsa–Romanowsky |
| WBCBench2026 Phase-1<sup>20, 21</sup> | External evaluation | 8,288 | 13 / 7 | PB | Wright–Giemsa, single site |
| TXL-PBC<sup>27</sup> | Detector training | 1,260 img | 3 / 1 | PB | 18,143 boxes |

Every training source was passed through the Stage-1 detector before use: the highest-confidence box was kept, and an image whose best box scored below 0.70 was excluded rather than cropped (the per-dataset pools of the leave-one-dataset-out analysis used source-specific thresholds). The filter removed 37.8% of the densely packed bone-marrow BMCMHF fields but far less of each single-cell peripheral-blood source; persource attrition is in Supplementary Section S1 and the harmonized counts in Table 1. For BMCMHF this is a deliberate purity-for-yield trade (Section 5).

After label harmonization and preprocessing, the merged corpus comprised 127,938 single cells across nine cell types. Acquisition magnification and on-specimen pixel scale are not held constant across sources —itself one facet of the acquisition shift studied here (Supplementary Section S1). Data were partitioned strictly at the *dataset* level for the external comparison (Section 2.1). The internal train/validation/test split, by contrast, was a per-class random 80/10/10 split at the *cell* level (a single fixed split, seed 42, held constant across the proposed model’s three seeds). Because most source datasets release no patient or slide identifiers, that split is not guaranteed to be patient-disjoint, so internal metrics may be optimistic. The leave-one-dataset-out classifier is held out at the dataset level in every fold, but the detector is not: a small detector exposure (≈3% of held-out cells) is confined to the Raabin-WBC and PBC folds, and the other three folds have none (Supplementary Section S1).

The internal label space comprised nine cell types (Neutrophil, Lymphocyte, Monocyte, Eosinophil, Basophil, Promyelocyte, Myeloblast, Lymphoblast, and Erythroblast). The merged corpus was markedly imbalanced (16.5:1 between the most and least frequent classes in the seven-class evaluation space), and its class distribution differed from the external set (Supplementary Table S2), which motivates macro-F1 as the primary endpoint and balanced accuracy as a secondary one. The datasets also differ visibly in staining, color balance, and background (Supplementary Fig. S1), which is the cross-dataset domain shift this study targets.

### 2.3 Label harmonization

Because the external benchmark does not provide the same fine-grained taxonomy, we mapped predictions and ground truth to a common seven-class space before computing external metrics. Blast group absorbs myeloblast, lymphoblast and erythroblast on the training side and the benchmark’s single “blast” label; neutrophil absorbs band and segmented neutrophils; lymphocyte absorbs variant lymphocytes and prolymphocytes; basophil, eosinophil, monocyte and promyelocyte map to themselves; and plasma cells, metamyelocytes and myelocytes have no counterpart in the evaluation space and are excluded. This mapping was applied identically to every model. The first-step crosswalk from each dataset’s native classes to the nine internal labels (and the native classes dropped) is given in Supplementary Table S3 (merged-corpus exceptions: Supplementary Section S1); lymphoblast training examples come from a single source (HRLS), and the abnormal promyelocytes characteristic of acute promyelocytic leukemia (APL) are not a distinct training category. Throughout, the seven evaluation classes are abbreviated basophil (BASO), eosinophil (EOS), lymphocyte (LYM), monocyte (MONO), neutrophil (NEU), promyelocyte (PROMY), and blast group (BLAST). The classifier retains nine output classes, but external BLAST metrics assess the merged category rather than discrimination among its constituent classes (Supplementary Table S4).

### 2.4 Detection-and-cropping stage

WBC detection used YOLO11-M^17,18^ trained on the 882-image training split of TXL-PBC^27^ (Table 1) for up to 150 epochs at 512 × 512 (batch 32, seed 3407; the automatic optimizer setting selected AdamW, learning rate 1.43 × 10^−3^ decaying linearly to 1.46 × 10^−4^); early stopping on the validation split (patience 30) ended training after epoch 137 (the best-epoch checkpoint, epoch 107, was retained). Training used WBC boxes only, the one class needed downstream (the checkpoint keeps the dataset’s three-class head, but only WBC has training instances). On the TXL-PBC validation split (252 images, 257 WBC boxes; Ultralytics 8.4.82 default validation settings at 512 px) it reached precision 0.996, recall 0.987, mAP@50 0.994 and mAP@50– 95 0.880. A three-class variant (adding red blood cells and platelets) localized WBCs comparably and was not retained (Supplementary Section S1).

At inference the detector is run at 512 px with a confidence threshold of 0.30 and an NMS IoU of 0.70; the highest-confidence box is expanded by 10% on each side and cropped, and the crop is square-padded, resized to 256 px and center-cropped to 224 before classification (Fig. 2). Coverage is reported with every pipeline result (Section 3.2).

**Fig. 2.**
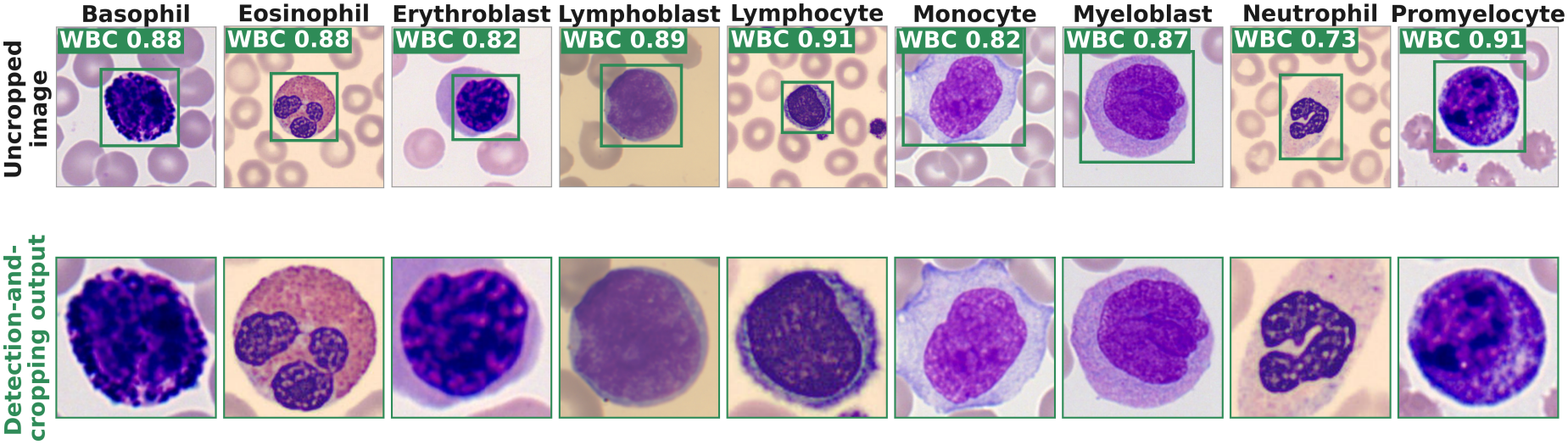
Stage 1, detection and cropping. Top: uncropped images for each of the nine training classes, overlaid with the bounding box predicted by the pipeline’s YOLO11 detector (green); the number in the “WBC” tag is the detector’s confidence (0–1) that the boxed cell is a white blood cell. Bottom: the resulting 224 × 224 crop of the box expanded by 10% on each side, which retains the surrounding background (the square padding applied before classification is not shown). Image sources as in Fig. 1.

### 2.5 Classification models

The primary classifier was `swin_base_patch4_window7_224`^19^ at 224 px, trained with class-weighted cross-entropy and morphology-preserving color augmentation (referred to below as the *proposed* model); the convolutional baselines were ResNet50,^28^ VGG16-BN^29^ and EfficientNet-B0.^30^ The proposed model and all three convolutional baselines were trained on one common corpus and partition (102,345 training, 12,803 held-out internal images, nine internal classes), so the class-weighted cross-entropy rows of Table 3b vary backbone and recipe but not the data. Retraining an earlier, smaller-corpus version of these baselines on this common corpus raised their external macro-F1 by 0.077–0.129. Recipes and seeds are in Supplementary Table S5, hardware in Section 2.9.

### 2.6 Pipeline routes and the detection-and-cropping ablation

To test whether the crop-trained classifier requires the detection-and-cropping stage, we compared the deployable pipeline (uncropped image →YOLO crop →Swin; Fig. 1) against the same classifier given the uncropped image (resized directly into the classifier). An off-the-shelf Cellpose^31, 32^ cropper was also run with and without the surrounding background (Supplementary Table S6).

### 2.7 Class imbalance and operating point

For every model trained with class-weighted cross-entropy (the proposed model, the other class-weighted Swin-B and ResNet50 configurations, the convolutional baselines and the fine-tuned DinoBloom-B), class imbalance was handled in the loss by weighting each class by the inverse square root of its training count, 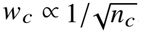, which up-weights rare classes without the extreme weights produced by inverse-frequency weighting. Every main result uses argmax decisions (Section 2.8). Because the minority classes (Promyelocyte, Blast group) are clinically important, we also ran a post hoc, secondary one-versus-rest threshold analysis for these two classes: each threshold maximized that class’s F1 on the internal held-out test split, used as a validation proxy, and was then applied unchanged to the external set (Supplementary Table S11).

### 2.8 Evaluation metrics and statistics

The primary endpoint was macro-F1 of the deployable full-pipeline route on the WBCBench2026 Phase-1 split, counting detection failures (*n* = 8,080; Section 2.1): a labeled image on which the detector returns no box receives a no-detection output that matches no class, so it adds a false negative to its true class, and macro-F1 is the unweighted mean of the seven class F1 scores (the no-detection output is not averaged as an eighth class). Pre-cropped comparisons (*n* = 8,061) are secondary. We additionally report accuracy, balanced accuracy, weighted F1, per-class precision/recall/F1 and minimum-class recall. Three seed aggregations are used and labeled wherever they appear: (i) the mean and population sd of per-seed metrics (a descriptive spread over three runs), used in the main-text tables, where for external results each seed’s nine-class argmax is mapped to the seven evaluation classes; (ii) a three-seed probability ensemble 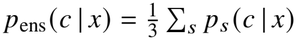, with the nine-class probabilities summed into the seven classes before the argmax, used for ensemble point estimates, bootstrap intervals, calibration, the confusion matrices and the error analysis; and (iii) a three-seed majority vote of the per-seed seven-class labels, used for the paired tests of Supplementary Section S7, which require one hard label per image (a three-way tie takes the seed-42 label). The two ensemble rules disagree on a few images (64 of 8,061 for the proposed model). Image-level 95% confidence intervals for each model’s ensemble macro-F1 used 2,000 nonparametric bootstrap replicates (percentile method). Paired comparisons used McNemar’s exact test on per-image correctness and a paired bootstrap of the macro-F1 difference (5,000 replicates; the recipe-selection bootstraps of Section 3.6 used 1,000 replicates on probability ensembles); equivalence was tested by two one-sided tests at a ±0.02 margin (set before the shared-protocol AdamW comparison was run; not a clinical threshold), i.e. established only if the 90% interval lies within the margin. The one seed-level comparison (proposed versus fine-tuned DinoBloom-B) is a paired *t*-test over the three seed-matched pairs. All image-level intervals treat images as independent and describe fixed fitted models, so they include neither seed variability nor patient clustering; the competition release of Phase-1 used here exposes no per-image patient identifier, so no patient-clustered bootstrap was run, and the intervals are anti-conservative for patient-level generalization. Cross-source variability is characterized separately by the leave-one-dataset-out analysis. All secondary comparisons are exploratory and were not adjusted for multiple testing. The route, sample size, decision rule and aggregation behind each main classification analysis are listed in Supplementary Section S17.

### 2.9 Implementation and reproducibility

The Swin-B, BPaCo and DinoBloom-B models, the frozen probe, full-pipeline detector inference and every WBCBench2026 evaluation were run, or reproduced image for image, with one software stack (Python 3.10, PyTorch 2.5.1, timm 1.0.27, Ultralytics 8.4.82); the convolutional baselines were trained, and the leave-one-dataset-out models evaluated, in a second (Python 3.12, PyTorch 2.9.0, timm 0.9.12); the detector was trained with Ultralytics 8.3.170, and Cellpose ran separately (versions in Supplementary Table S5). Training used NVIDIA RTX 4090 (24 GB), RTX 4060 Ti (16 GB) and RTX A6000 (48 GB) GPUs. The proposed model and the class-weighted-CE convolutional baselines were initialized from timm’s default pretrained weights, whose pretraining data differ by backbone (Swin-B: ImageNet-22k, fine-tuned on ImageNet-1k; ResNet50 and VGG16-BN: ImageNet-1k; EfficientNet-B0: Noisy Student weights, trained on ImageNet-1k with unlabeled JFT-300M images), and were trained at 224 × 224 with class-weighted cross-entropy and ImageNet normalization. The other rows of Table 3 depart from this: the BPaCo rows started from the same timm weights but used unweighted cross-entropy plus the BPaCo contrastive term; the fine-tuned DinoBloom-B was initialized from DinoBloom’s self-supervised weights and fine-tuned with the proposed recipe’s hyperparameters, with its layer-wise learning-rate decay applied per transformer block rather than per Swin block group (a group of transformer blocks operating at one spatial resolution) and with timm’s default drop-path settings (configured maximum rate 0.1 for Swin-B, 0 for the ViT); and the frozen DinoBloom-B probe kept those weights fixed and fitted an unweighted logistic-regression classifier (Supplementary Section S7). The proposed model’s augmentation is deliberately constrained — square padding and a mild color jitter with no geometric, blur, or noise augmentation, unlike generic strong-augmentation policies such as RandAugment^33^ — so that cell morphology is preserved. The convolutional neural network (CNN) baselines differ from it in augmentation and optimization settings but use the same imbalance handling (class-weighted cross-entropy, no resampling, no mixup). For WBCBench2026 evaluation all models used one deterministic transform (square padding, resize to 256 px, i.e. 1.14×, center-crop to 224) with horizontal-flip test-time augmentation, and every route in Table 2 and Supplementary Table S6 was scored with the same evaluator, as were the Phase-2 results of the deployable pipeline (Section 3.2), those first scored otherwise being re-evaluated post hoc; the resolution sweep and the missed-versus-detected comparison keep their own settings (Supplementary Section S17). Code for the inference pipeline, together with training scripts and environment specifications, is publicly available (Code availability).

**Table 2.**
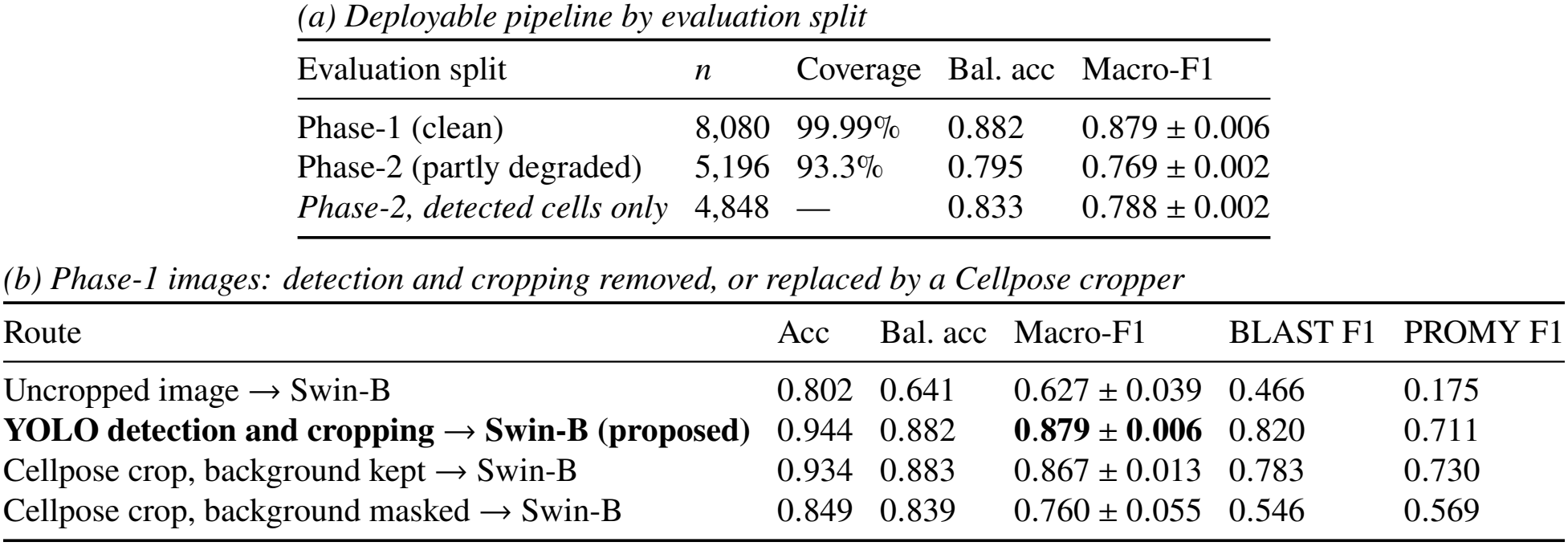
External performance of the deployable pipeline and the effect of the detection-and-cropping stage on WBCBench2026 (seven classes; mean ± sd over seeds 42/62/82; columns without sd are seed means). (a) The same deployable pipeline (uncropped image →YOLO →Swin-B) on the clean Phase-1 and the partly degraded Phase-2 split, in which about 60% of images were synthetically degraded by the organizers; images on which the detector found no cell count as errors, “coverage” is the fraction of images with a detection, and the final row is the optimistic detected-cells-only view. (b) The same crop-trained Swin-B checkpoints on the Phase-1 images (*n* = 8,080) with or without detection and cropping, or with an off-the-shelf Cellpose cropper that keeps or masks the surrounding background (Supplementary Table S6). The pre-cropped classifier route (0.863 ± 0.028, *n* = 8,061) differs in detector threshold, crop settings and *n*, so its difference from the YOLO row is not the effect of the detection-and-cropping stage.

| *(a) Deployable pipeline by evaluation split* |  |  |  |  |  |
|---|---|---|---|---|---|
| Evaluation split | *n* | Coverage | Bal. acc | Macro-F1 |  |
| Phase-1 (clean) | 8,080 | 99.99% | 0.882 | 0.879 ± 0.006 |  |
| Phase-2 (partly degraded) | 5,196 | 93.3% | 0.795 | 0.769 ± 0.002 |  |
| *Phase-2, detected cells only* | 4,848 | — | 0.833 | 0.788 ± 0.002 |  |
| *(b) Phase-1 images: detection and cropping removed, or replaced by a Cellpose cropper* |  |  |  |  |  |
| Route | Acc | Bal. acc | Macro-F1 | BLAST F1 | PROMY F1 |
| Uncropped image → Swin-B | 0.802 | 0.641 | 0.627 ± 0.039 | 0.466 | 0.175 |
| **YOLO detection and cropping → Swin-B (proposed)** | 0.944 | 0.882 | **0.879** ± **0.006** | 0.820 | 0.711 |
| Cellpose crop, background kept → Swin-B | 0.934 | 0.883 | 0.867 ± 0.013 | 0.783 | 0.730 |
| Cellpose crop, background masked → Swin-B | 0.849 | 0.839 | 0.760 ± 0.055 | 0.546 | 0.569 |

## 3 Results

Three Phase-1 macro-F1 figures recur for the proposed system: reporting views of the same detector and Swin checkpoint set, not three independently trained systems. The primary endpoint, 0.879, is the coverage-aware full-pipeline evaluation of uncropped images (seed mean, *n* = 8,080; Table 2); 0.863 is the pre-cropped classifier route (seed mean, *n* = 8,061; Table 3); and 0.893 is the three-seed probability ensemble of that pre-cropped route (Supplementary Tables S8 and S9). Across the full-pipeline endpoint and two complementary classifier-level summaries, external macro-F1 remained within 0.863–0.893, so the main result did not depend on a single input route or seed-aggregation rule, supporting the stability of the full pipeline.

**Table 3.**
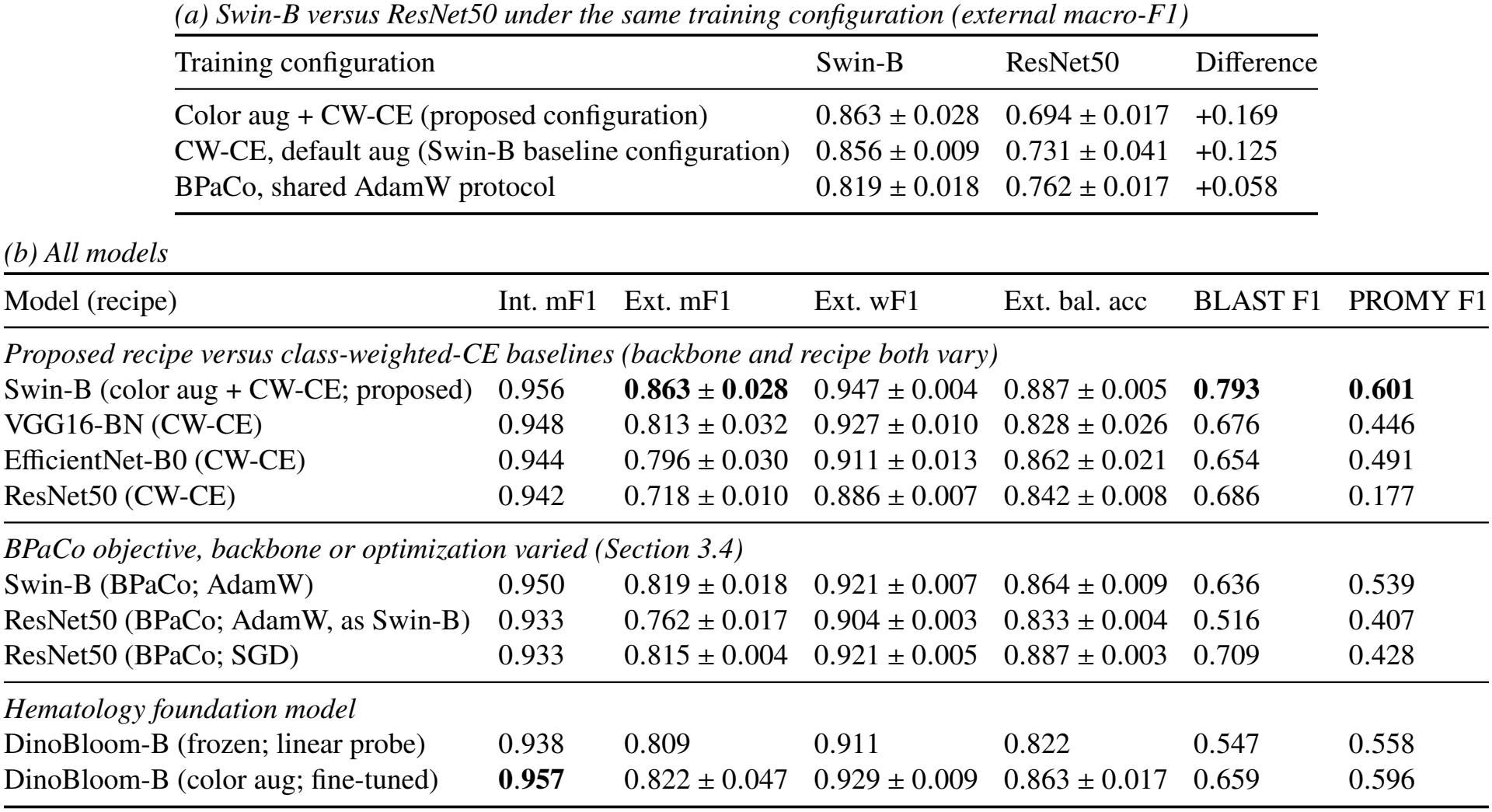
Backbone, training configuration and representation on the external benchmark, evaluated on pre-cropped WBCBench2026 Phase-1 cells (Section 2.1; seven classes, horizontal-flip test-time augmentation, *n* = 8,061). Values are mean ± sd over three seeds; columns without sd are seed means, and the frozen linear probe is a single fit. (a) Swin-B and ResNet50 trained under the same configuration (Section 3.4); differences are computed from unrounded seed means, and per-seed values are in Supplementary Section S7. (b) All models; every row shares one corpus and partition, and internal macro-F1 is nine-class on the same held-out set (*n* = 12,803). The two AdamW BPaCo rows share one training protocol and differ in their backbone-specific model configuration; the SGD row is an earlier arm with backbone-specific optimization (Supplementary Section S7). **Bold** marks the highest internal and the highest external macro-F1, which fall on different rows (Section 3.3), and the highest BLAST and PROMY F1. Int./Ext. mF1, internal/external macro-F1; wF1, weighted F1; bal. acc, balanced accuracy; BLAST, blast group; PROMY, promyelocyte; CW-CE, class-weighted cross-entropy; aug, augmentation.

| *(a) Swin-B versus ResNet50 under the same training configuration (external macro-F1)* |  |  |  |  |  |  |
|---|---|---|---|---|---|---|
| Training configuration | Swin-B | ResNet50 | Difference |  |  |  |
| Color aug + CW-CE (proposed configuration) | 0.863 ± 0.028 | 0.694 ± 0.017 | +0.169 |  |  |  |
| CW-CE, default aug (Swin-B baseline configuration) | 0.856 ± 0.009 | 0.731 ± 0.041 | +0.125 |  |  |  |
| BPaCo, shared AdamW protocol | 0.819 ± 0.018 | 0.762 ± 0.017 | +0.058 |  |  |  |
| *(b) All models* |  |  |  |  |  |  |
| Model (recipe) | Int. mF1 | Ext. mF1 | Ext. wF1 | Ext. bal. acc | BLAST F1 | PROMY F1 |
| *Proposed recipe versus class-weighted-CE baselines (backbone and recipe both vary)* |  |  |  |  |  |  |
| Swin-B (color aug + CW-CE; proposed) | 0.956 | **0.863** ± **0.028** | 0.947 ± 0.004 | 0.887 ± 0.005 | **0.793** | **0.601** |
| VGG16-BN (CW-CE) | 0.948 | 0.813 ± 0.032 | 0.927 ± 0.010 | 0.828 ± 0.026 | 0.676 | 0.446 |
| EfficientNet-B0 (CW-CE) | 0.944 | 0.796 ± 0.030 | 0.911 ± 0.013 | 0.862 ± 0.021 | 0.654 | 0.491 |
| ResNet50 (CW-CE) | 0.942 | 0.718 ± 0.010 | 0.886 ± 0.007 | 0.842 ± 0.008 | 0.686 | 0.177 |
| *BPaCo objective, backbone or optimization varied (Section 3.4)* |  |  |  |  |  |  |
| Swin-B (BPaCo; AdamW) | 0.950 | 0.819 ± 0.018 | 0.921 ± 0.007 | 0.864 ± 0.009 | 0.636 | 0.539 |
| ResNet50 (BPaCo; AdamW, as Swin-B) | 0.933 | 0.762 ± 0.017 | 0.904 ± 0.003 | 0.833 ± 0.004 | 0.516 | 0.407 |
| ResNet50 (BPaCo; SGD) | 0.933 | 0.815 ± 0.004 | 0.921 ± 0.005 | 0.887 ± 0.003 | 0.709 | 0.428 |
| *Hematology foundation model* |  |  |  |  |  |  |
| DinoBloom-B (frozen; linear probe) | 0.938 | 0.809 | 0.911 | 0.822 | 0.547 | 0.558 |
| DinoBloom-B (color aug; fine-tuned) | **0.957** | 0.822 ± 0.047 | 0.929 ± 0.009 | 0.863 ± 0.017 | 0.659 | 0.596 |

### 3.1 The crop-trained classifier requires detection and context-preserving cropping

Detection and cropping were necessary at inference for the crop-trained classifier: giving it the uncropped image collapsed external macro-F1 to 0.627 ± 0.039, driven by large-scale Blast group over-prediction and loss of minority classes; the trained detection-and-cropping stage restored macro-F1 to 0.879 ± 0.006 with the smaller across-seed sd (Table 2b). The effect is one of train/test consistency — the detector supplies the in-distribution single-cell input the crop-trained classifier expects, with near-complete coverage on this clean split (8,079 / 8,080). The requirement is for a context-preserving crop rather than for this detector in particular: an off-the-shelf Cellpose cropper that keeps the surrounding background reached a comparable 0.867 ± 0.013, while the same cropper with the background masked out fell to 0.760 ± 0.055 — the classifier depends on the surrounding background, which the masking removes (Table 2b; Supplementary Table S6).

### 3.2 The Phase-2 shift degrades both pipeline stages

We evaluated the same deployable pipeline on the WBCBench2026 Phase-2 split, whose scanner, blur, and illumination degradation is applied by the organizers.^21^ Performance dropped at both stages. First, detection failed far more often: on the harmonized-label subset, detector coverage fell from 8,079 /8,080 (99.99%) on the clean Phase-1 split to 4,848 /5,196 (93.3%) on Phase-2; the 348 undetected images count as errors. Second, classification of the cells that were still found was also worse. Together, external macro-F1 fell from 0.879 ± 0.006 to 0.769 ± 0.002 and balanced accuracy from 0.882 to 0.795 (Table 2a) — consistent across seeds, but larger than the detected-cells-only view would suggest (macro-F1 0.788 ± 0.002 if the 348 detection failures are discarded). Passed to the classifier under identical input treatment, the cells the detector missed were substantially harder than a class-matched control of detected cells (accuracy 0.727 versus 0.862; Supplementary Table S12), so the detector’s misses marked a harder-to-classify subset; 73% of them would still have been labeled correctly.

### 3.3 Internal macro-F1 did not calibrate external performance

The four class-weighted cross-entropy models (the proposed Swin-B, VGG16-BN, EfficientNet-B0 and ResNet50) were trained on the same corpus and partition and reached high, tightly clustered internal macro-F1 on one identical held-out set (0.942–0.956; across-seed sd ≤0.001). Externally the same models spread more than ten times as widely (0.718–0.863), led by the proposed Swin (Table 3b). Among these four models the internal column ranks them exactly as the external column does, but internal macro-F1 does not calibrate the external *level*: a 0.014 internal spread accompanied a 0.145 external one. The ordering fails as well, but only once the training configuration or the representation changes: the highest internal score in the table (DinoBloom-B fine-tuned; below) is not the best external model, and the same ResNet50 scored 0.009 lower internally but 0.097 higher externally when trained with BPaCo and its own SGD optimization (below). Both observations hold with the internal scores mapped to the seven evaluation classes (Supplementary Section S17). Contrasting weighted-F1 with macro-F1 clarifies the shortfall: externally the CNN baselines retained a high weighted-F1 (0.886–0.927), which is carried by the frequent Neutrophil and Lymphocyte classes, while their macro-F1 fell further because performance on the minority classes dropped — Promyelocyte F1 from 0.601 (Swin) down to 0.177 (ResNet50), and Blast group F1 from 0.793 to 0.65–0.69 (Table 3). The proposed model’s advantage over the class-weighted-CE baselines is thus concentrated where it matters clinically, in the blast and promyelocyte categories; Section 3.4 compares the backbones with more of the configuration held fixed. Per-class F1 rests on few cells for the smallest external classes (Promyelocyte *n* = 42, Basophil *n* = 74; Supplementary Table S2; per-class F1 by backbone: Supplementary Table S10). For macro-F1, bootstrapping the three-seed ensembles over external images (Methods) gave non-overlapping 95% confidence intervals — Swin [0.893 0.874, 0.911]; CNN ensembles 0.751–0.848, upper bounds ≤0.865 — so the advantage exceeds the image-level sampling variability of these fixed ensembles (per-model intervals in Supplementary Table S9).

### 3.4 Swin-B outperformed ResNet50 under all three training configurations

Three comparisons hold more of the configuration fixed (Table 3a). First, the proposed color-augmentation configuration run on ResNet50 (same corpus, three seeds) gave 0.694 ± 0.017 externally — *below* that back-bone’s class-weighted-CE baseline (0.718) — against 0.863 for Swin-B under the same configuration, a seed-mean gap of 0.169; with SGD (no warmup, layer-wise decay or mixed precision; Supplementary Section S7) instead of AdamW, ResNet50 reached 0.727 ± 0.020. The configuration’s layer-wise learning-rate decay is defined over Swin’s four block groups and gives ResNet50 a single backbone learning rate. Second, we trained Swin-B and ResNet50 with BPaCo^34, 35^ (three seeds each) under one shared training protocol — the same objective, augmentation, AdamW optimizer, learning rate, weight decay, schedule and mixed precision — so that the two arms differ only in their backbone-specific model configuration (architecture, pretrained weights and timm’s default drop-path setting; Table 3; hyperparameters in Supplementary Section S7). The ResNet50 arm of this comparison was added post hoc, after an earlier BPaCo pair was found to have used backbone-specific optimizers. Under the shared protocol Swin-B remained ahead (0.819 ± 0.018 versus 0.762 ± 0.017; higher on every seed), and the three-seed majority-vote ensembles differed by +0.062 macro-F1 (95% CI [+0.044, +0.079]; McNemar *p* < 0.001), an interval lying wholly above the ±0.02 equivalence margin (Supplementary Table S7). Third, the configuration of the Swin-B class-weighted-CE baseline (default augmentation; Section 3.6), applied to both backbones, gave 0.856 ± 0.009 (Swin-B) versus 0.731±0.041 (ResNet50; the 0.718 ResNet50 row of Table 3 used the CNN baselines’ own recipe, Supplementary Table S5). Swin-B was thus higher in all three comparisons, on every seed, by 0.058 (BPaCo; unrounded seed means) to 0.169.

ResNet50 was also sensitive to its optimization settings. Its earlier BPaCo arm (SGD, full precision) reached 0.815 ± 0.004 externally, 0.053 above the shared-protocol arm, and came close to Swin-B’s BPaCo arm (0.819; Supplementary Section S7); as it differs from the AdamW arms in optimizer, learning rate, weight decay and precision, it is not one of the three comparisons.

We also asked whether external robustness merely reflects representation quality by comparing against a published hematology foundation model, DinoBloom-B,^36^ a DINOv2 vision transformer self-supervised on ∼380,000 single-cell blood and marrow images. Under a frozen linear probe its external macro-F1 was 0.809: within the range of the class-weighted-CE convolutional baselines (0.718–0.813) and below the proposed Swin (0.863). Fully fine-tuned with the proposed recipe’s hyperparameters (Section 2.9), it reached an internal macro-F1 of 0.957 — marginally *above* the proposed Swin’s 0.956 — yet an external macro-F1 of 0.822 ± 0.047, below the proposed model on every matched seed and with a larger across-seed spread (Table 3; per-seed values in Supplementary Section S7). On the shared external images the proposed ensemble outperformed DinoBloom’s (McNemar *p* < 0.001; Supplementary Table S7); the per-seed difference was not significant at three seeds (paired *p* ≈0.15). The comparison is confounded in one direction: DinoBloom’s pretraining corpus includes three of the five training sources used here (AML-LMU, Raabin-WBC and 80% of BMCMHF), which favors it. Both models were run at 224 px, which is also the resolution at which DinoBloom was self-supervised.^36^ Under leave-one-dataset-out the direction reverses — fine-tuned DinoBloom-B exceeds the proposed model on every fold — with the same confound on the AML-LMU, Raabin-WBC and BMCMHF folds; on the PBC and HRLS folds, absent from its pretraining corpus,^36^ it still led, by 0.009 and 0.018 (Supplementary Section S16).

### 3.5 Leave-one-dataset-out reveals the non-redundant value of source diversity

To test whether the remaining sources can replace the morphology, acquisition and class variation of an absent dataset, we performed classifier-level LODO evaluation across the five training datasets’ pools (Table 1). Because a held-out dataset contains only a subset of the nine classes, we report macro-F1 over the classes actually present (support> 0; full nine-class values in Supplementary Note 1). Present-class macro-F1 ranged from 0.59 (AML-LMU) to 0.91 (PBC), mean 0.72 (Table 4). The internal ∼0.95 is not a like-for-like comparator (nine-class macro-F1 on the pooled internal split, versus present-class scores on each held-out dataset’s own cells), so the folds are read as a *relative* measure of how well the other four sources substituted for each absent one. Image count alone did not ensure substitution: with AML-LMU held out, the training set still held 2,027 HRLS myeloblasts, yet AML-LMU Myeloblast recall was 0.025 (F1 0.048), most read as Lymphocyte —a missed blast. HRLS is the only Lymphoblast source, so its fold had no Lymphoblast training support

**Table 4.** Classifier-level leave-one-dataset-out generalization (mean ± sd, seeds 42/52/62; Supplementary Note 1), on each held-out dataset’s pool of cells that passed the detector filter, without test-time augmentation. As a classifier-level evaluation it does not count detection failures, unlike the deployable-pipeline endpoint (Table 2); the detector saw about 3% of the PBC and Raabin-WBC held-out cells (Section 2.2). Present-class macro-F1 averages only classes present in the held-out dataset (full-label-space values: Supplementary Note 1); columns without sd are seed means. LODO models use the proposed recipe, trained on the other four sources’ training splits and checkpoint-selected on their validation splits. **Bold** highlights the present-class macro-F1 column, its best held-out value (PBC), and its across-dataset mean. and Lymphoblast F1 0.000 in all three seeds of both LODO arms, the proposed-recipe arm and an earlier mixup/CutMix arm^37, 38^ (read as Myeloblast — a zero-shot class; Supplementary Section S14 and Note 1; Section 4). BMCMHF, the only bone-marrow source, scored 0.606 ± 0.030, the widest seed spread (per-seed values in Supplementary Note 1), consistent with a non-substitutable bone-marrow domain, but holding it out also removed nearly all Promyelocyte and most Erythroblast training support (Supplementary Table S15), so its gap cannot be attributed to bone-marrow morphology alone. Per-class values: Supplementary Section S14.

| Held-out | # classes | Accuracy | **Macro-F1 (present)** | Bal. acc (present) |
|---|---|---|---|---|
| AML-LMU | 8 | 0.782 | 0.590 ± 0.008 | 0.718 |
| PBC | 6 | 0.919 | **0.909 ± 0.015** | 0.888 |
| Raabin-WBC | 5 | 0.844 | 0.742 ± 0.015 | 0.716 |
| HRLS | 7 | 0.778 | 0.763 ± 0.008 | 0.808 |
| BMCMHF | 7 | 0.702 | 0.606 ± 0.030 | 0.643 |
| Mean |  |  | **0.722** | 0.754 |

### 3.6 Color augmentation: best macro-F1, lower minimum recall

On pre-cropped external cells (*n* = 8,061), three Swin-B recipes were compared as three-seed probability ensembles (Supplementary Table S8): the color-augmentation recipe, a class-weighted-CE baseline with default augmentation, and an aggressive recipe combining mixup^37^ with BPaCo and classifier re-training (cRT^39^). The color-augmentation ensemble reached the highest macro-F1 (0.893) and the highest Blast group and Promyelocyte F1 (0.841 and 0.702, versus 0.829 and 0.687 for the baseline and 0.822 and 0.611 for the aggressive recipe, which was lowest on both). A paired image-level bootstrap run during recipe selection gave macro-F1 differences of +0.013 (95% CI [+0.0004, +0.025]) for the color-augmentation ensemble over the class-weighted-CE baseline (0.893 versus 0.881) and +0.030 ([+0.016, +0.044]) over the aggressive recipe (0.864); as the recipe was selected on these images (Section 2.1), the intervals are not confirmatory. The class-weighted-CE baseline had the highest minimum-class recall (0.810 versus 0.786) and balanced accuracy (0.913 versus 0.908). In the threshold analysis (Methods), thresholds tuned on the internal test split raised external Promyelocyte F1 slightly (0.702 to 0.719) while Blast group F1 changed little (0.841 to 0.839); even thresholds tuned on the external set itself would add little (Supplementary Table S11). The ensemble’s external calibration error was low (expected calibration error 0.017, Brier score 0.074; Supplementary Fig. S2), with a mild overconfidence (0.970 mean confidence versus 0.954 accuracy), mostly in the top confidence bin (90% of predictions; 0.998 versus 0.984).

### 3.7 Naive resolution up-scaling collapses performance

The 224-trained model was not robust to naive inference-only resolution scaling (the same weights applied to crops square-padded, resized and center-cropped to 256 or 384 px, with the model’s patch grid scaled to match): on the pre-cropped set (*n* = 8,061), macro-F1 fell from 0.863 ± 0.027 at the native 224 px (the sweep’s own evaluator; Supplementary Section S17) to 0.830 ± 0.039 at 256 px and collapsed to 0.464 ± 0.084 at 384 px, consistently across all three seeds. At 384 px every class degraded, the four smallest classes most on average (across seeds, Blast group F1 → 0.07–0.52 and Promyelocyte F1 → 0.00–0.35).

### 3.8 Errors concentrate on morphologically similar classes

Internally, per-class recall was uniformly high (0.90–0.98; Fig. 3), and in the nine-class space the three constituent blast types were also well separated (Supplementary Table S4). Externally, recall was preserved for Neutrophil (0.99), Eosinophil (0.97) and Lymphocyte (0.94) but fell for the two clinically important minority categories: Blast group recall dropped from 0.95 to 0.84 (with 11% called Lymphocyte), and Promyelocyte from 0.96 to 0.79 (with 10% called Basophil and 7% Monocyte) — so external misclassifications concentrated among morphologically similar cell types. In a structured breakdown of the 368 external errors (4.6% of 8,061 predictions), the most frequent were the bidirectional Blast group ↔Lymphocyte confusion (65 and 64 errors; 35% of all errors) and Lymphocyte ↔ Monocyte (63 and 27; 24%) — a cluster of mononuclear and immature cells; only 3% were confusions between the mature granulocyte classes. Promyelocyte errors were dominated by false alarms (19 versus 9 false negatives) at its very low external prevalence of 0.5%.

**Fig. 3.**
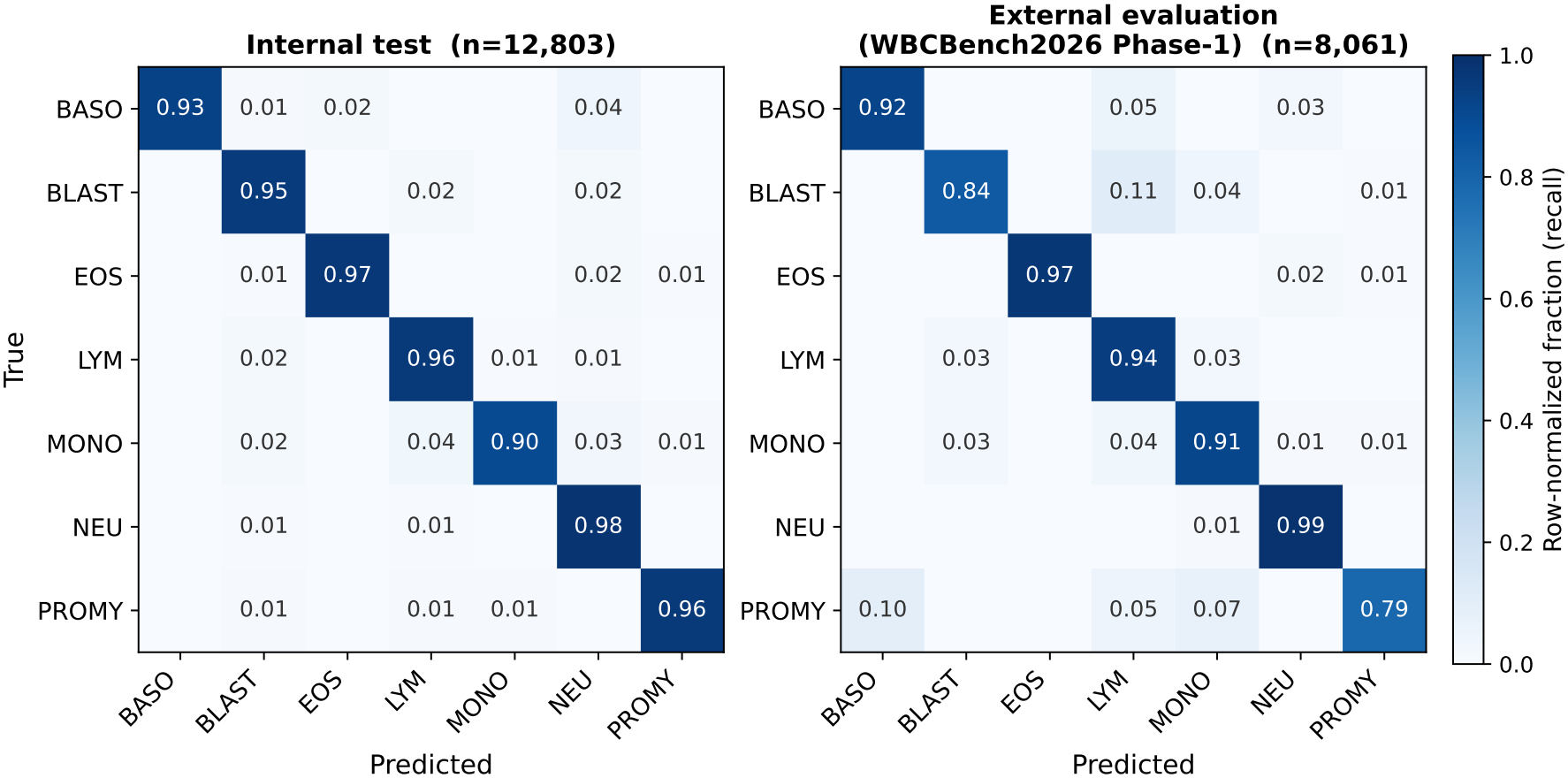
Row-normalized confusion matrices (recall) of the proposed Swin model (three-seed probability ensemble, seven-class mapped label space) on the internal test set (left, *n* = 12,803) and the external WBCBench2026 Phase-1 pre-cropped set (Section 2.1; right, *n* = 8,061). Entries below 0.005 are not annotated, and rows may not sum to 1 because of rounding. BASO, basophil; BLAST, blast group; EOS, eosinophil; LYM, lymphocyte; MONO, monocyte; NEU, neutrophil; PROMY, promyelocyte.

### 3.9 Two-hematologist review of hardest-class disagreements

Two board-certified hematologists at different hospitals reviewed all 205 external cells on which the proposed model (pre-cropped route, three-seed probability ensemble) and the WBCBench2026 label disagreed and either the label or the prediction was Blast group or Promyelocyte. They rated the same crops independently, without discussing them, as *model right* (benchmark label wrong, model correct), *model wrong* (benchmark label right) or *ambiguous*; the review form showed each crop with the benchmark label and the model prediction (Supplementary Section S13).

The two hematologists gave the same verdict on 73 of the 205 crops (35.6%): both rated the model right on 21 cells (10.2%), both rated it wrong on 43 (21.0%), and both rated 9 (4.4%) ambiguous. At least one rated 22 (10.7%) ambiguous, and on 119 (58.0%) one rated the model right and the other wrong (Supplementary Table S13). All benchmark metrics stand unchanged.

## 4 Discussion

Across training sources that differ in acquisition and framing (Supplementary Section S1), the detection- and-cropping stage gave the classifier cell-focused inputs that keep the surrounding background, which the crop-trained classifier required (Table 2b). Combined with multi-source training and morphology-preserving color augmentation, the pipeline reached an external macro-F1 of 0.879 on an unseen benchmark site and 0.769 on the partly degraded Phase-2 split. The paragraphs below consider what drove the external spread, what the detection-and-cropping stage contributes, what the leave-one-dataset-out results imply for training data, which minority-class trade-offs remain, what the expert review implies for benchmark labels, and what these findings mean for deployment.

### Internal metrics and backbone choice

Across the three training configurations evaluated on both back-bones, Swin-B achieved higher external macro-F1 than ResNet50 (+0.058 to +0.169 on pre-cropped cells; Section 3.4), including under the shared AdamW–BPaCo protocol, supporting a consistent backbone-associated external advantage of Swin-B in this task. ResNet50’s own external score moved by 0.053 between two BPaCo optimization settings while its internal macro-F1 remained nearly unchanged. Internal macro-F1 ranked the four class-weighted-CE models as the external benchmark did but did not calibrate their external level, and changing the training configuration or representation inverted the ordering (Section 3.3). These results qualify the use of internal performance as a proxy for external performance, a relationship studied in the “accuracy on the line” literature;^10, 12^ prior work distinguishes robustness gains under distribution shift from those expected from higher in-distribution accuracy.^11^ Backbone and training configuration should therefore be chosen jointly.

### Detection and cropping are part of the model, not preprocessing

Removing the detector lowered external macro-F1 more (0.879 →0.627) than the entire backbone-and-recipe spread of Table 3 (0.169), and on the Phase-2 split the detector failed alongside the classifier (Section 3.2). A crop-trained classifier and its detector are therefore one system with one operating point, and classifier accuracy measured on curated crops describes a component rather than a deployable pipeline.

### Representing each cell class across multiple sources

The LODO findings support representing the same cell classes across multiple data sources when developing robust WBC classifiers. For example, the presence of myeloblast training examples from HRLS did not ensure successful transfer to AML-LMU, showing that class representation alone does not guarantee cross-source generalization. The held-out sources contained morphology, acquisition and class variation that was not fully covered by the remaining sources, in keeping with documented cross-site degradation;^6, 8^ LODO did not compare source diversity with image count at a fixed total. Lymphoblasts, which come from HRLS alone (Section 2.3), are the limiting case: the HRLS fold was zero-shot rather than domain-shifted, so the internal myeloblast–lymphoblast separation (Supplementary Table S4) has no source-disjoint test. Neither LODO nor the external metrics, which merge all three blast types, validate the myeloblast-versus-lymphoblast (AML versus ALL) distinction. Fine-tuned DinoBloom-B exceeded the proposed model on every LODO fold but not on WBCBench2026 (Section 3.4); as its pretraining corpus includes three of the five held-out sources, neither result establishes a general robustness advantage.

### Minority-class gains and remaining trade-offs

The proposed recipe’s gains over both comparator Swin-B recipes (Section 3.6) extended to the underrepresented Blast group and Promyelocyte classes rather than being confined to the dominant classes. Morphology-preserving color augmentation is the class of intervention shown to mitigate stain variation across laboratories in computational pathology;^7^ here it was more effective than mixup with BPaCo and cRT, which did not improve minority-class F1, while its margin over the Swin-B class-weighted-CE baseline was small (95% interval just above zero; Section 3.6). As that baseline also used class weighting, the gains reflect the recipe rather than class weighting, and trade-offs remain: the baseline kept the higher minimum-class recall, and Promyelocyte F1 remained the main limitation. The ensemble’s external Blast group recall of 0.84 (Section 3.8) means that 16% of benchmark-labeled blasts were assigned to a non-blast class. Because the blast group merges erythroblasts (nucleated red-cell precursors) with leukemic myelo- and lymphoblasts, the external BLAST metric is a broad immature-cell score, not a leukemia-specific blast count; and because APL promyelocytes were not a separate training category (Section 2.3), the promyelocyte output is a normal/reactive flag rather than APL screening. A deployment prioritizing blast sensitivity over precision could adopt a recall-weighted blast operating point — a lower one-versus-rest blast threshold —set by a sensitivity-target sweep on held-out internal data rather than on the external benchmark.

### Benchmark labels as ground truth

External scores treat the benchmark labels as ground truth. Those labels were assigned by five experienced hematopathologists, with ambiguous cases discussed in consensus sessions.^21^ When two further hematologists independently reviewed the 205 hardest-class disagreements, they agreed on 73 crops: on 43 both judged the model wrong, on 9 both rated the cell ambiguous, and on 21 both judged the model correct and the benchmark label wrong (Section 3.9). Even expert-assigned labels on a public benchmark can therefore contain errors, and a second expert reading can reach a different label, as reported for manual differentials^1^ and for immature and blast-like cells.^3^ Studies that score models against such benchmarks should keep in mind that the benchmark label taken as ground truth may itself be wrong.

### Pathology-informatics relevance

Two elements bear on deployment. First, the coverage-aware protocol reports the operating point a triage queue would see; because Phase-2 detection failures concentrated on harder cells (Section 3.2), a no-detection is a usable signal for routing cells to human review. Second, the internal–external dissociation argues for external, coverage-aware acceptance testing before a WBC classifier is trusted in a new laboratory information system.

## 5 Limitations

1. **Data coverage.** WBCBench2026 evaluates transfer to a new acquisition site with standardized staining, but does not by itself establish performance across multiple sites or staining protocols. LODO provides complementary cross-source evidence at the classifier level. Phase-1 and Phase-2 also involve different patients, so cohort differences may contribute to the observed Phase-2 decrease. The Phase-2 degradation is synthetic and benchmark-defined, so it measures robustness to a controlled stressor rather than to real artifacts such as out-of-focus regions, smear thickness or staining drift.
2. **Evaluation design.** Because augmentation and test-time augmentation were chosen on Phase-1 (Section 2.1), the Phase-1 estimates for the selected recipe are optimistically selected, whereas Phase-2 played no part in these choices.
3. **Scope of the detection-and-cropping ablation.** A classifier trained directly on uncropped images was not evaluated, so the ablation does not compare two-stage processing with alternatives trained on uncropped images.
4. **Scope of the model comparisons.** The comparisons evaluate backbone-specific model configurations rather than architecture alone. Pretrained weights and settings such as drop-path differ, and the shared AdamW–BPaCo comparison between Swin-B and ResNet50 was added post hoc. The comparators include CNN baselines and DinoBloom-B evaluated through both a frozen linear probe and full fine-tuning. No head-to-head comparison with the challenge-organizer baseline was performed because official test labels were absent from the competition files used here.
5. **Detector-based selection of the development data.** Of the BMCMHF images passed to the detector, 37.8% were excluded at the 0.70 confidence threshold (Supplementary Section S1). Internal metrics therefore describe detector-selected cells and do not cover the excluded images.
6. **Computational performance.** Inference latency, throughput and memory use were not measured. The pipeline combines approximately 20 million detector parameters with 87 million classifier parameters.
7. **Images used for the expert review.** The hematologists rated the pre-cropped cells that the classifier receives, cut without margin (Supplementary Sections S6 and S13), rather than the uncropped images. Only a few crops cut off part of the cell, but such a crop can make the cell impossible to rate; the crops also lack the surrounding red cells that give a size reference, and the highest-confidence box need not be the benchmark’s target cell. Having the experts read the uncropped images would avoid this.

*Future work* should extend evaluation of the complete pipeline to multiple institutions and acquisition conditions. To investigate the contribution of source diversity more directly, controlled training comparisons should hold total image counts and per-class counts fixed while varying the number of sources represented within each class. Further robustness studies should target localization and classification under challenging image conditions and assess native 256 / 384-px training rather than inference-only resizing. Additional priorities include measuring inference latency, throughput and memory use.

## 6 Conclusion

This study developed a two-stage YOLO11–Swin-B pipeline for cross-dataset white blood cell classification, combining WBC detection and context-preserving cropping with multi-source classifier training and morphology-preserving color augmentation. The full pipeline achieved an external macro-F1 of 0.879 ± 0.006 on WBCBench2026 Phase-1. Detection and context-preserving cropping were essential for maintaining the performance of the crop-trained classifier, while the proposed color-augmentation recipe achieved higher macro-F1 and higher Blast group and Promyelocyte F1 than the comparator recipes in the Swin-B ensemble analysis. On pre-cropped cells, Swin-B consistently outperformed ResNet50 across all three evaluated training configurations. The LODO findings further support representing the same cell classes across multiple data sources when developing robust classifiers. Together, these results support the proposed two-stage design and emphasize that robust WBC classification requires attention to both model configuration and the diversity of the training data.

## Ethics statement

This study used only publicly available, previously published, de-identified image datasets, each in accordance with its license and data-use terms; no new human-subject data were collected and no patient-identifiable information was accessed. Institutional review board approval was therefore not required.

## Data availability

All datasets are public and cited in Table 1: AML-Cytomorphology_LMU (TCIA, doi:10.7937/tcia.2019.36f5o9ld; CC BY 3.0); Bone-Marrow-Cytomorphology_MLL_Helmholtz_Fraunhofer (TCIA, doi:10.7937/TCIA.AXH3-T579; CC BY 4.0); PBC (Mendeley Data, doi:10.17632/snkd93bnjr.1; CC BY 4.0); Raabin-WBC^25^ (https://raabindata.com; free access with citation under the Raabin Health Database terms); HRLS (figshare, doi:10.6084/m9.figshare.22680517; CC0 1.0); TXL-PBC (figshare, doi:10.6084/m9.figshare.27073186; CC BY 4.0); and WBCBench2026 (Kaggle competition wbc-bench-2026, listed as CC BY-NC-SA 4.0; we used the competition files downloaded in November 2025, whose rules restrict redistribution; a later Hugging Face release corrects 257 labels, removes 10 images and adds a patient hash). Example images in Figs. 1 and 2 and Supplementary Fig. S1 are reproduced only from PBC, HRLS, AML-LMU and BMCMHF under the licenses above; Raabin-WBC and WBCBench2026 images are not reproduced. The aggregate and per-class metric tables underlying the figures and the evaluation code are available in the code repository; WBCBench2026 images, labels and per-image predictions are not redistributed.

## Code availability

The two-stage inference pipeline, training scripts, configuration files, evaluation code, and environment specifications (Supplementary Table S5) for the complete workflow are publicly available at https://github.com/NTUBIME34/wbc-crossdataset-code. The trained detector and classifier checkpoints (three seeds each for the proposed model and for the BPaCo comparisons) will be released upon publication.

## Declaration of generative AI and AI-assisted technologies in the manuscript preparation process

During the preparation of this work, the authors used AI-assisted writing tools (Claude Opus 4.8, Anthropic) to improve the language and readability of portions of the manuscript. After using these tools, the authors reviewed and edited the content as needed and take full responsibility for the content of the publication, and confirm that no AI tool is listed as an author or meets authorship criteria. No generative-AI tool was used to create or alter any figure, image, or scientific result reported in this work.

## CRediT authorship contribution statement

**Che-Wei Chang**: Conceptualization, Methodology, Software, Formal analysis, Investigation, Data curation, Visualization, Writing – original draft. **Kuo-Jui Sun**: Validation (clinical/morphological review of model classifications), Writing – review & editing. **Cheng-Hong Tsai**: Validation (clinical/morphological review of model classifications), Writing – review & editing. **Chien-Yu Chen**: Conceptualization, Supervision, Writing – review & editing.

## Appendix A. Supplementary data

Supplementary material (Sections S1–S17 and Supplementary Note 1) contains all per-class, per-dataset, provenance, training-recipe and analysis-specification detail.

