## Supplementary Material for "A Two-Stage YOLO–Swin Transformer Pipeline for Cross-Dataset White Blood Cell Classification"

Unless stated otherwise, all external results are on the WBCBench2026 Phase-1 pre-cropped evaluated set ($n$ = 8,061) in the harmonized seven-class space, reported as mean ± population standard deviation over seeds 42/62/82 (convolutional baselines: 42/43/44, Table S5); the routes of Section S6 that start from the uncropped image use all 8,080 labeled images and Section S12 uses Phase-2. Class abbreviations: basophil (BASO), eosinophil (EOS), lymphocyte (LYM), monocyte (MONO), neutrophil (NEU), promyelocyte (PROMY), blast group (BLAST).

### S1. Dataset provenance, preprocessing, and appearance variation

**Detector filtering of the training sources.** Every training source was passed through the Stage-1 detector before use, keeping the highest-confidence box and discarding any image whose best box scored below a confidence threshold: 0.30 (0.70 for BMCMHF) in the per-dataset pools used by the leave-one-dataset-out analysis (Tables S1 and S15), and 0.70 for every source in the merged internal corpus ("Merged internal corpus" below). The four peripheral-blood sources are distributed as single-cell crops and lose almost nothing at 0.30. BMCMHF, the only bone-marrow source, has densely packed fields and loses 37.8%; of its 92,007 retained cells, 79,880 fall in nine categories after label harmonization — the seven internal labels it maps to plus myelocytes and metamyelocytes, which have no internal counterpart — and the 73,096 in the seven internal labels form its harmonized pool (Table S15). The Raabin-WBC row is its released training split, the only Raabin split used in this study.

Table S1
Detector filtering of each training source for its per-dataset pool, at a confidence threshold of 0.30 (0.70 for BMCMHF). "Released" is the number of images as distributed, except for PBC, AML-LMU and BMCMHF, where it counts the images passed to the detector (PBC's immature-granulocyte and platelet images, AML-LMU's 15 smudge-cell images and BMCMHF's artifact, not-identifiable, other-cell and smudge-cell categories were not passed to it); "excluded" images were dropped rather than cropped at low confidence; all were detection failures except 12 of the 16 HRLS images, which could not be decoded in the downloaded copy.

| Source | Released | Excluded | Retained | % excluded |
|---|---|---|---|---|
| AML-LMU | 18,350 | 12 | 18,338 | 0.07 |
| PBC | 11,849 | 4 | 11,845 | 0.03 |
| Raabin-WBC | 10,175 | 21 | 10,154 | 0.21 |
| HRLS | 16,027 | 16 | 16,011 | 0.10 |
| BMCMHF | 147,869 | 55,862 | 92,007 | 37.78 |

**Detector variant that was not used.** A three-class detector variant (WBC, red blood cells, platelets) was trained on the same images and scored against the identical WBC ground truth. It performed comparably on the WBC class itself (precision 0.996, recall 0.996, mAP@50 0.994, mAP@50–95 0.894) — adding the erythrocyte and platelet labels changed WBC localization only marginally (+0.009 recall and +0.014 mAP@50–95, against unchanged precision and mAP@50). The WBC-only model was retained for simplicity; the three-class variant was not evaluated in the full pipeline.

**External attrition chain.** From the 8,288 released Phase-1 uncropped images, cropping at a detector confidence of 0.70 yields 8,269 crops; the 19 images whose top detection falls below the threshold are dropped (13 lymphocyte,

6 segmented neutrophil). Removing the 208 cells (2.5%) whose native labels have no counterpart in the nine-class output taxonomy (myelocyte 108, metamyelocyte 98, plasma cell 2) leaves the evaluated pre-cropped set of $n = 8{,}061$. The full-pipeline route skips the 0.70 cropping step: it starts from all 8,288 fields at a detector confidence of 0.30, removes the same 208 taxonomy-unmatched fields and evaluates the remaining $n = 8{,}080$, counting the one field with no detection as an error rather than dropping it. The Phase-2 split is handled the same way: of its 5,350 released images, 154 carry a native label with no counterpart in the output taxonomy and are excluded before any metric is computed, leaving the $n = 5{,}196$ over which coverage and the coverage-aware macro-F1 are reported. The detector returns no box for 357 of the 5,350; 348 of those fall inside the labeled subset and are the ones counted as errors.

**Merged internal corpus.** The five harmonized single-cell pools sum to 128,628 cells; the merged corpus used for internal training and evaluation contains 127,938. The difference was quantified image by image (SHA-256; decoded pixels for the JPEG-sourced BMCMHF and Raabin-WBC images). The merged corpus, cropped separately at 0.70, lacks the 966 pool images whose best box scored between 0.30 and 0.70 (0.8%, mostly Neutrophil and Monocyte; AML-LMU 599, PBC 66, Raabin-WBC 155, HRLS 146, BMCMHF 0) and contains 276 images from native classes outside the crosswalk of Table S3 — 103+99 AML-LMU/HRLS band neutrophils under Neutrophil, 50 BMCMHF immature lymphocytes under Lymphocyte and 24 AML-LMU monoblasts under Myeloblast (0.2% and 0.4% of those classes), the last two being label noise. Fifteen byte-identical duplicate pairs remain, each within a single partition, so no identical file is shared between the training, validation, and test splits; 158 images share a perceptual hash across splits. The leave-one-dataset-out experiments are built directly from the per-dataset pools (Table S15) and contain none of the 276.

**Overlap between detector and classifier training sources.** TXL-PBC, used only to train the detector, is itself an integration of four public sources: Raabin-WBC (500 images), PBC (500), BCCD (160) and BCDD (100). Two of these — PBC and Raabin-WBC — also appear among the classifier training datasets, so a subset of those images is shared between detector- and classifier-training. We quantified that overlap exactly: TXL-PBC's distribution records the source dataset and original filename of every image, and the detector was trained on its 882-image training split (composition: Raabin-WBC 358, PBC 353, BCCD 101, BCDD 70). Matching those original filenames against the leave-one-dataset-out held-out sets gives 358 of the 10,154 held-out Raabin-WBC cells (3.53%) and 353 of the 11,845 held-out PBC cells (2.98%). The exposure is confined to the detection-and-cropping stage, and the AML-LMU, HRLS and BMCMHF folds have none, because none of these sources contributes to TXL-PBC. All leave-one-dataset-out arms score the same detector crops of the same cells.

WBCBench2026 is not one of TXL-PBC's four sources and postdates it. We also checked this directly: the SHA-256 content hash of every one of the 1,260 TXL-PBC images was compared against the hash of every one of the 8,288 WBCBench2026 Phase-1 fields. All 1,260 and all 8,288 are distinct within their own sets, and the two sets share no hash and no filename, so not one detector-training image reaches the external benchmark. The comparison is rerun from the two hash lists (`analysis/evidence/leakage_audit/`) by `analysis/verify_tables.py`; we keep hashes rather than images because the competition rules under which WBCBench2026 was obtained restrict redistribution of its data.

**Acquisition magnification and pixel scale.** Source acquisition magnifications were ×100 oil immersion for the peripheral-blood datasets (AML-LMU, Raabin-WBC, HRLS) and ×40 for the bone-marrow dataset (BMCMHF), while the PBC images were acquired on a CellaVision DM96 analyzer (objective not reported by the source). Two sources report an on-specimen scale: the AML-LMU TCIA collection record (Matek et al., 2019) gives a manufacturer-specified 14.14 pixels/$\mu$m ($\approx$ 0.071 $\mu$m/px) and HRLS reports $\approx$ 42 pixels/$\mu$m ($\approx$ 0.024 $\mu$m/px), an $\approx 3\times$ difference; the remaining sources (Raabin-WBC, BMCMHF, PBC, and WBCBench2026) do not report a usable physical scale. Because all crops are resized to 224 px for the classifier, physical pixel scale is not held constant across sources.

**Appearance variation across the sources.** Fig. S1 shows the same three cell types in four of the five training sources.

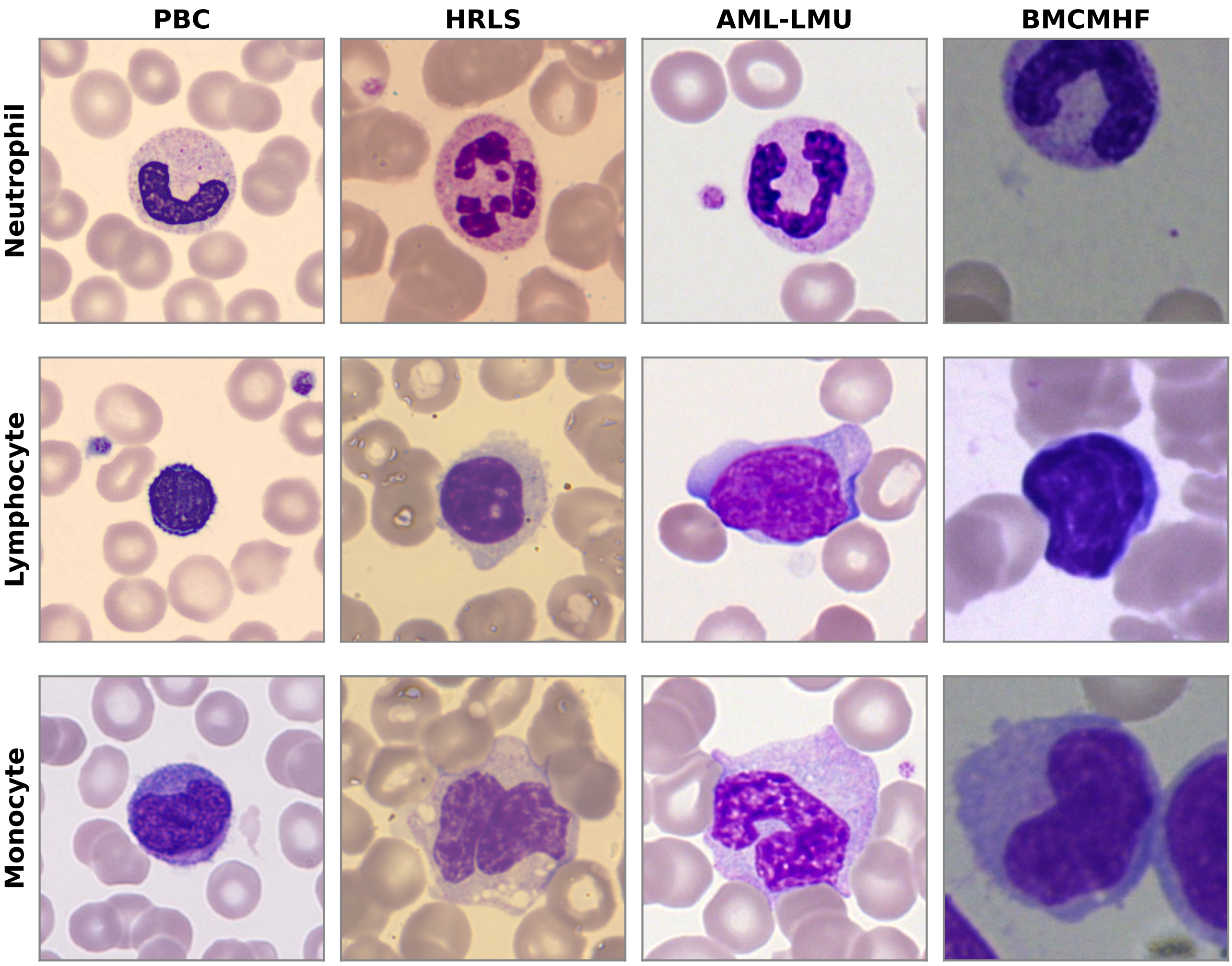


Fig. S1. Domain shift across the source datasets. The same cell types (rows: neutrophil, lymphocyte, monocyte) are shown for four of the five training datasets (columns), illustrating differences in staining, color balance, and background that constitute cross-dataset domain shift. Images are the dataset-original files as distributed (no detector cropping applied), reproduced at a common display size. Example cells are reproduced from PBC (Acevedo et al., 2020), HRLS (Bodzas et al., 2023), AML-LMU (Matek et al., 2019), and BMCMHF (Matek et al., 2021), each under its own public license (CC BY 4.0, CC0 1.0, CC BY 3.0, and CC BY 4.0, respectively; see Data availability). Raabin-WBC, also used for model training, is omitted here because its images are not licensed for reproduction.

### S2. Class distribution of the internal and external sets

Table S2
Class distribution of the merged internal corpus (training, validation and test splits) and the external WBCBench2026 Phase-1 set in the harmonized seven-class space. The external column is the pre-cropped evaluated set ($n$ = 8,061; see the "External attrition chain" paragraph of Section S1 for the attrition from the released split). The internal corpus is imbalanced (16.5:1 most- to least-frequent classes), and the external set is shifted toward neutrophils, lymphocytes and monocytes and away from Blast group and Promyelocyte.

| | Internal corpus | | External | |
|---|---|---|---|---|
| Class | $n$ | % | $n$ | % |
| Neutrophil | 47,551 | 37.2 | 4,018 | 49.8 |
| Lymphocyte | 30,811 | 24.1 | 2,433 | 30.2 |
| Blast group | 23,422 | 18.3 | 572 | 7.1 |
| Eosinophil | 8,766 | 6.9 | 181 | 2.2 |
| Monocyte | 8,526 | 6.7 | 741 | 9.2 |
| Promyelocyte | 5,988 | 4.7 | 42 | 0.5 |
| Basophil | 2,874 | 2.2 | 74 | 0.9 |
| Total | 127,938 | | 8,061 | |

### S3. Per-dataset label crosswalk (native class → internal nine-class label)

Table S3 gives the label crosswalk used to build the five harmonized single-cell pools (Table S15); the leave-one-dataset-out analysis uses these pools directly. Only explicitly mapped native classes were kept (e.g. band and segmented neutrophils → Neutrophil for BMCMHF); all other native classes were dropped. Two consequences matter for interpretation. The BMCMHF "Blast" native class (~12,000 images) was dropped rather than remapped, so no dataset contributes an unspecified "blast" to the internal Myeloblast/Lymphoblast labels; those two labels come only from AML-LMU (myeloblast) and HRLS (myeloblast, lymphoblast). The HRLS "Normoblast" class was not mapped to Erythroblast and was therefore excluded, so HRLS contributes no Erythroblast cells.

Table S3
First-step label crosswalk. For each dataset, each internal nine-class label is listed with the native source class(es) that map into it; the final column lists native classes that were dropped. Band neutrophils were merged into Neutrophil for BMCMHF but excluded from the AML-LMU and HRLS pools ($n$ = 109 and 99); the merged corpus departs from this crosswalk in 276 images (Section S1).

| Dataset | Internal label ← native class(es) | Dropped native classes |
|---|---|---|
| AML-LMU | Basophil←BAS; Eosinophil←EOS; Erythroblast←EBO; Lymphocyte←LYT, LYA; Monocyte←MON; Myeloblast←MYO; Neutrophil←NGS; Promyelocyte←PMO | KSC (smudge cell), MMZ (metamyelocyte), MYB (myelocyte), MOB (monoblast), NGB (band neutrophil, $n$ = 109), PMB (bilobed promyelocyte, $n$ = 18) |
| BMCMHF | Basophil, Eosinophil, Erythroblast, Lymphocyte, Monocyte, Promyelocyte ← same; Neutrophil←Segmented + Band neutrophil | Blast, Abnormal eosinophil, Faggot cell, Hairy cell, Immature lymphocyte, Metamyelocyte, Myelocyte, Plasma cell, Proerythroblast; Artifact, Not identifiable, Other cell, Smudge cell (not passed to the detector; Table S1) |
| PBC | Basophil, Eosinophil, Erythroblast, Lymphocyte, Monocyte, Neutrophil ← same (lower-cased native names) | ig (immature granulocytes), platelet |
| Raabin-WBC | Basophil, Eosinophil, Lymphocyte, Monocyte, Neutrophil ← same | (none) |
| HRLS | Basophil, Eosinophil, Lymphoblast, Lymphocyte, Monocyte, Myeloblast ← same; Neutrophil←Neutrophil_seg | Normoblast (nucleated red-cell precursor), Neutrophil_band ($n$ = 99) |

Mapping verified against each dataset's native folders and the built nine-class training folders (content-hash checks for the ambiguous BMCMHF and AML-LMU sub-classes). AML-LMU codes follow the dataset's own `abbreviations.txt`.

### S4. Internal discrimination of blast sub-types

For external evaluation, myeloblast, lymphoblast, and erythroblast are collapsed into a single BLAST class because the external benchmark provides only a coarse "blast" label (see Label harmonization, main text). This collapse is a property of the benchmark, not of the model: on the internal nine-class test set the classifier separates the three sub-types with high per-class F1 and almost never confuses them with one another (Table S4). Because myeloblasts are associated with AML and lymphoblasts with ALL, this separation would matter clinically, but it has no source-disjoint support (Section S14; main-text Discussion).

Table S4
Internal nine-class per-class performance for the three blast sub-types (three-seed mean ± sd), and their mutual confusion.

| Sub-type | Clinical meaning | $n$ | F1 | Confused with other blast types |
|---|---|---|---|---|
| Myeloblast | AML blast | 582 | 0.974 ± 0.001 | 0.2% |
| Lymphoblast | ALL blast | 257 | 0.992 ± 0.002 | 1.4% |
| Erythroblast | nucleated red precursor | 1,506 | 0.924 ± 0.001 | 0.0% |

Source: internal nine-class test reports (`test_results.json`) and confusion matrices for seeds 42/62/82. Mutual-confusion is the fraction of each true sub-type assigned to either of the other two blast sub-types.

### S5. Training recipe registry

This table lists the proposed model and the class-weighted-CE convolutional baselines; the BPaCo and foundation-model configurations (Section S7), the augmentation and imbalance ablation (Sections S7, S8 and S17), the leave-one-dataset-out models (Supplementary Note 1) and the detector (main Methods) are specified where they are reported. Training resolution, normalization and external evaluation are as in the main Methods; the internal held-out scores of these classifiers use square padding, resize to 1.1× (246 px) and center-crop to 224, without test-time augmentation.

Table S5
Training recipes of the proposed model and the convolutional baselines (the class-weighted-CE reference rows in panel b of main-text Table 3); Section S7 reports comparisons that hold more of the training configuration fixed.

| | Proposed Swin-B | CNN baselines |
|---|---|---|
| Backbone | `swin_base_patch4_window7_224` | ResNet50, VGG16-BN, EfficientNet-B0 |
| Initialization | timm `ms_in22k_ft_in1k` (ImageNet-22k, fine-tuned on ImageNet-1k) | timm defaults: ResNet50 `a1_in1k` and VGG16-BN `tv_in1k` (ImageNet-1k); EfficientNet-B0 `tf_efficientnet_b0.ns_jft_in1k` (Noisy Student: ImageNet-1k plus unlabeled JFT-300M) |
| Optimizer | AdamW, lr $10^{-4}$, wd $10^{-4}$ (biases and normalization weights exempt) | AdamW, lr $10^{-4}$, wd $10^{-4}$ |
| Schedule | 5-epoch warmup, then cosine decay; layer-wise lr decay 0.8; head lr ×2 | cosine-annealing decay |
| Batch / epochs | 128 / 30 | 64 / 30 |
| Augmentation | square padding, resize 1.08×, center-crop 224, color jitter (brightness, contrast, saturation 0.12; hue 0.03); no geometric, blur, or noise augmentation | square padding, resize 1.1×, random-resized-crop 224 (scale 0.8–1.0), random horizontal flip, color jitter (brightness, contrast, saturation 0.2; hue 0.05) |
| Imbalance handling | class-weighted cross-entropy; no resampling, no mixup | class-weighted cross-entropy; no resampling, no mixup |
| Seeds | 42/62/82 | 42/43/44 for each convolutional baseline |
| Precision | automatic mixed precision, gradient checkpointing | full precision (FP32) |

Environments. Swin-B, BPaCo, DinoBloom-B and Section S7 ResNet50 training, the frozen probe, full-pipeline detector inference and all WBCBench2026 evaluation: Python 3.10.20, PyTorch 2.5.1 (CUDA 12.1, cuDNN 9.1), torchvision 0.20.1, timm 1.0.27, Ultralytics YOLO 8.4.82, scikit-learn 1.7.2, NumPy 2.2.6 (`journal_env.yml`). Convolutional-baseline training and leave-one-dataset-out evaluation: Python 3.12.12, PyTorch 2.9.0 (CUDA 12.8), torchvision 0.24.0, timm 0.9.12. Detector training: Ultralytics 8.3.170; the versions behind the detector crops (per-dataset pools and pre-cropped set, November 2025; merged corpus, March 2026) were not recorded. Cellpose 4.1.1 (`cpsam` model) in a separate environment. GPUs: NVIDIA RTX 4090 (24 GB), RTX 4060 Ti (16 GB) and RTX A6000 (48 GB).

### S6. Cropper comparison: trained detector versus off-the-shelf segmentation

Beyond the background effect reported in the main text (route D versus route C), an off-the-shelf segmentation cropper is *competitive but not superior* to the trained detector (C 0.867 versus B 0.879), and re-cropping the route C crops with the trained YOLO detector (route E) gives the highest point estimate (0.882 ± 0.019 versus 0.879 ± 0.006 for route B). Route B, the deployable pipeline defined in the main text, predates this comparison, which used the external labels (Section S17); it was retained.

Table S6
Cropping routes on the external benchmark, all scored with the same classifier evaluator (main Methods). "Uncropped" passes the whole image (368×368 or 368×370 px) to that evaluator without detection or cropping. Route A is a classifier-only reference: the pre-cropped set ($n$ = 8,061) holds the same detector's highest-confidence box at threshold 0.70, cropped without margin, so its difference from route B is not the effect of the detection-and-cropping stage. Route B crops the box at threshold 0.30 with a 10% margin; routes C and D take a 224 × 224 window centered on the Cellpose mask nearest the image center (a mask was found in every image), D whitening every pixel outside that mask; route E runs the detector on each route C crop and re-crops its top box with a 5% margin (every crop yielded a box). The other rows use all 8,080 labeled images, route B counting its one image without a detection as an error. Columns without sd are seed means; bold marks the deployable route.

| Route | Description | Acc | Bal. acc | Macro-F1 | BLAST F1 | PROMY F1 |
|---|---|---|---|---|---|---|
| — | Uncropped image → Swin (no detection or cropping) | 0.802 | 0.641 | 0.627 ± 0.039 | 0.466 | 0.175 |
| A | Pre-cropped → Swin (classifier reference) | 0.946 | 0.887 | 0.863 ± 0.028 | 0.793 | 0.601 |
| B | Image → **YOLO** → Swin (deployable) | 0.944 | 0.882 | **0.879 ± 0.006** | 0.820 | 0.711 |
| C | Image → Cellpose (keep background) → Swin | 0.934 | 0.883 | 0.867 ± 0.013 | 0.783 | 0.730 |
| D | Image → Cellpose (background removed) → Swin | 0.849 | 0.839 | 0.760 ± 0.055 | 0.546 | 0.569 |
| E | Image → Cellpose crop → YOLO re-crop → Swin | 0.951 | 0.899 | 0.882 ± 0.019 | 0.817 | 0.680 |

Source: unified re-evaluation of the three checkpoints with one evaluator (Section S17). Cellpose v4.1.1 with the Cellpose-SAM model (Stringer et al., 2021; Pachitariu et al., 2025).

### S7. Backbone comparisons across training configurations; foundation models

The class-weighted-CE reference rows in panel b of main-text Table 3 contrast the proposed Swin model against class-weighted cross-entropy CNN baselines, so they confound the backbone with the training configuration (Table S5). The comparisons below (panel a of main-text Table 3) hold more of the configuration fixed (three seeds 42/62/82 each; external WBCBench2026 Phase-1, $n$ = 8,061). Three training configurations were evaluated on both backbones, in the order of the main text: class-weighted cross-entropy with color augmentation, balanced parametric contrastive learning (BPaCo) under a shared protocol, and class-weighted cross-entropy with default augmentation; ResNet50 was also trained with SGD under the first two. In the second, shared-protocol comparison, Swin-B and ResNet50 were trained with identical training arguments — objective, augmentation, AdamW optimizer, learning rate, weight decay, schedule and mixed precision — and differ only in their backbone-specific model configuration (architecture, pretrained weights and timm's default drop-path setting); this ResNet50 arm was added post hoc, after the earlier ResNet50 BPaCo arm was found to have used SGD with backbone-specific settings. Swin-B led externally (0.819±0.018 versus 0.762±0.017; per seed 0.836/0.827/0.795 versus 0.741/0.761/0.783). In the majority-vote ensembles its advantage was concentrated in the Blast group, Eosinophil and Promyelocyte classes (F1 differences +0.122, +0.114 and +0.182; Promyelocyte support 42), and ResNet50 was higher only for Monocyte (−0.022). The earlier SGD arm reached 0.815 ± 0.004, 0.053 above the shared-protocol ResNet50 arm in seed means, an effect of the whole optimization bundle (optimizer, learning rate, weight decay and precision). Under the first, color-augmentation configuration of Table S5 (Swin-B: the proposed model, 0.863), ResNet50 reached 0.694 ± 0.017 externally (per seed 0.672/0.713/0.697; internal 0.931), and 0.727 ± 0.020 (0.707/0.721/0.755; internal 0.939) with the SGD settings of the earlier BPaCo arm, which also omit warmup, layer-wise decay and mixed precision. The third configuration, class-weighted cross-entropy with the default augmentation of the Swin-B baseline in Table S8, gave 0.856±0.009 (Swin-B) and 0.731±0.041 (ResNet50) externally (per seed 0.849/0.869/0.849 versus 0.678/0.777/0.738; internal 0.959 versus 0.941). In both class-weighted configurations the AdamW ResNet50 arm was added post hoc and trained on the same split, in the same software environment and with the Swin-B arm's recorded hyperparameters except gradient checkpointing (a memory-saving setting, on for Swin-B only). Swin-B was therefore higher in all three configurations (by 0.169 under color augmentation, 0.058 under shared-protocol BPaCo and 0.125 under default augmentation, in seed means), and within each configuration every Swin-B seed exceeded every ResNet50 seed. In the class-weighted pairs the layer-wise learning-rate decay, defined over Swin's block groups, gives ResNet50 a single backbone learning rate (0.8× the base rate; head ×2).

Table S7

Paired comparisons on the shared 8,061 external images. The paired tests use three-seed majority-vote ensembles (main Methods; a three-way tie takes the seed-42 label), which, unlike probability ensembles, can be formed for the BPaCo runs, whose probabilities were not stored. McNemar's exact test is on per-image correctness; the paired bootstrap (5,000 replicates) resamples the macro-F1 difference Δ (first model minus second); TOST (two one-sided tests) establishes equivalence at a ±0.02 margin only if the 90% interval lies within it.

| Comparison | McNemar $p$ | Δ | 95% CI | 90% CI | TOST (±0.02) |
|---|---|---|---|---|---|
| Swin-B vs. ResNet50, BPaCo, shared AdamW protocol | <0.001 | +0.062 | [+0.044, +0.079] | [+0.047, +0.076] | not equivalent |
| Swin-B (AdamW) vs. ResNet50 (SGD), BPaCo | 0.21 | +0.010 | [−0.006, +0.026] | [−0.003, +0.023] | not established |
| Proposed vs. DinoBloom-B (fine-tuned) | <0.001 | +0.031 | [+0.015, +0.046] | [+0.018, +0.044] | not established |

Point estimates and ± sd for every row are in main-text Table 3. With a different seed breaking three-way ties, the first two Δ stay within 0.003 of the values shown, but the DinoBloom-B Δ ranges from +0.017 to +0.031 (every 95% interval excludes zero); its tie-free probability ensembles give +0.029 ([+0.016, +0.043]). BPaCo, balanced parametric contrastive learning.

**Interpretation of the paired tests.** For the earlier SGD arm no image-level difference was detected (McNemar $p = 0.21$; the 95% interval includes zero), but equivalence was not established either (the 90% interval reaches +0.023); the seed means differ by 0.005 (0.8195 versus 0.8148), and Swin-B was higher on two of the three seed-matched pairs (+0.016, +0.017, −0.019; differences of unrounded values, here and below). Against the SGD variant of the color configuration, by contrast, Swin-B remained higher on every seed (+0.179, +0.158, +0.069). The proposed model is better than the fine-tuned DinoBloom at the image level, even though the three-seed paired comparison is not significant ($p \approx 0.15$; seed-paired differences +0.051, +0.006 and +0.064).

**BPaCo hyperparameters.** All three BPaCo arms were trained for 30 epochs with batch size 128 at 224×224 with cosine decay and no warmup or layer-wise decay, from timm's default ImageNet-pretrained weights (Table S5), on the same nine-class training split (seed 42). The Swin-B arm and the shared-protocol ResNet50 arm used AdamW (learning rate $1 \times 10^{-4}$, weight decay 0.05) with automatic mixed precision and gradient checkpointing; the earlier ResNet50 arm used SGD (momentum 0.9, learning rate 0.02, weight decay $5 \times 10^{-4}$) in full precision. The training objective augments unweighted cross-entropy with a balanced parametric contrastive term, $\mathcal{L} = \mathcal{L}_{CE} + 0.25\,\mathcal{L}_{BPaCo}$ (Cai et al., 2024, which extends parametric contrastive learning, PaCo; Cui et al., 2021). The contrastive branch used a MoCo-style memory queue of size 8,192, a 128-dimensional projection through an MLP head, temperature 0.2, and PaCo-branch loss weights $(\alpha, \beta, \gamma) = (0.05, 1.0, 1.0)$. Each image yields two augmented views: the first applies a random-resized crop (scale 0.08–1), horizontal flip, color jitter (brightness, contrast and saturation 0.4) and RandAugment ($n = 2$ operations, magnitude $m = 10$ jittered with standard deviation 0.5, each operation applied with $p = 0.5$, as implemented in timm; Cubuk et al., 2020); the second applies a random-resized crop, the same color jitter, random grayscale ($p = 0.2$), Gaussian blur (kernel 3, $p = 0.5$) and horizontal flip. Mixup and CutMix were disabled. Internal held-out scores use resize to 256 px and center-crop to 224 without padding or test-time augmentation; external scores use the common external protocol. Exported checkpoints are plain classifier weights (the contrastive projection head is discarded at inference). The separate aggressive-recipe ablation (Table S8) instead adds to the configuration of the class-weighted-CE Swin-B baseline a BPaCo term (weight 0.1), mixup ($\alpha = 0.1$, no CutMix) and eight epochs of classifier re-training (cRT; Kang et al., 2020), which re-fits only the final linear classifier under class-balanced sampling with the backbone frozen.

**Foundation-model baseline.** DinoBloom-B (a DINOv2 ViT-B/14 self-supervised on ~380,000 single-cell blood and bone-marrow images; Koch et al., 2024) is used as a frozen linear-probe comparator: the model (timm `1aurent/vit_base_patch14_224.dinobloom`, Apache-2.0; the Hugging Face revision was not pinned at run time and is attributed retrospectively to the only locally cached snapshot, `4f2a9d8593a7d12efe210778ea856086 3fcacd03`) is run at 224 × 224 with the external protocol's preprocessing (square-pad, resize to 1.14×, center-crop to 224) on every split, and its 768-dimensional class-token embeddings are frozen, standardized on the training split, and classified by a multinomial logistic-regression probe (L-BFGS) fitted on the training split only, with no refit on training plus validation data. The inverse regularization strength ($C \in \{0.01, 0.1, 1, 10, 100\}$) and the class weighting (unweighted; inverse-square-root, $w_c \propto n_c^{-1/2}$ scaled to a mean sample weight of one; or inverse-frequency,

$w_c = n/(Kn_c)$, where $n_c$ is the training count of class $c$, $n$ the number of training images and $K = 9$) were selected by nine-class macro-F1 on the internal validation split, under a rule fixed before the full-data run. The selected probe was unweighted with $C$=0.01, the smallest value in the grid (the strongest regularization). Internal macro-F1 is computed on the common held-out set without test-time augmentation; externally, the probe's class probabilities for the original and horizontally flipped crop are averaged before the seven-class mapping. At their own validation-selected $C$ (also 0.01), the inverse-square-root and inverse-frequency probes reached external macro-F1 of 0.789 and 0.757 versus 0.809 unweighted, although their external balanced accuracy was higher (0.833 and 0.838 versus 0.822 for the unweighted probe). Summary internal-test and external metrics (macro-F1, weighted F1 and balanced accuracy) for all fifteen grid cells were computed in the same run and retained in `grid.csv`; selection used the validation column only. An unplanned post-run check below the grid edge (validation split only) raised validation macro-F1 by at most 0.0002; it was not used to alter the selected probe.

**Fully fine-tuned foundation model.** To test whether full fine-tuning changes the comparison with the proposed model, we additionally fine-tuned the entire backbone end-to-end under the proposed model's recipe (the proposed recipe of Table S5 — no contrastive term, the layer-wise decay adapted to the transformer's per-block depth; three seeds 42/62/82), a comparison of backbone-specific model configurations relative to the proposed Swin (differences listed in the main Methods). The released DinoBloom checkpoint stores 224 × 224 position embeddings (257 tokens), the resolution at which DinoBloom was itself self-supervised (Koch et al., 2024), so it was run at 224 × 224 with no position-embedding interpolation, under the pipeline's resolution and the identical external protocol. The `timm` configuration inherited from DINOv2 advertises 518 px; that is the base model's resolution, not DinoBloom's. The fine-tuned model reached an internal macro-F1 of 0.957 — marginally above the proposed Swin's 0.956, on the same corpus, partition and 12,803 held-out cells. The frozen probe's internal macro-F1 of 0.938 was computed with the external protocol's preprocessing (square padding, resize to 256 px, i.e. 1.14×, center-crop to 224), whereas the fine-tuned model's internal evaluation resized to 246 px (1.1×) before the same crop; neither used test-time augmentation. Only the external evaluation is identical for the two. Externally, the selected frozen probe yielded a macro-F1 of 0.809, whereas fine-tuning yielded 0.822 ± 0.047 across three seeds (per-seed 0.835/0.873/0.759), 0.013 above the probe on average, with one seed below it. The fine-tuned model remained below the proposed model's seed mean of 0.863 and below the proposed model on every matched seed (0.886/0.878/0.824 for seeds 42/62/82). Its across-seed spread was also larger (±0.047 versus ±0.028). Under leave-one-dataset-out the direction reverses (Table S16).

### S8. Augmentation and imbalance recipes on the external benchmark

Table S8
Augmentation / imbalance recipes on the external benchmark (pre-cropped classifier route, $n$ = 8,061; three-seed probability ensembles, horizontal-flip TTA). The three rows are all Swin-B; the class-weighted-CE row uses the default augmentation (random-resized crop, horizontal flip and color jitter) without the proposed color-only policy. **Bold** marks the proposed model's row and the best value in the macro-F1 and minimum-recall columns.

| Model (ensemble) | Acc | Bal. acc | Macro-F1 | Min recall | BLAST F1 | PROMY F1 |
|---|---|---|---|---|---|---|
| **Color aug (proposed)** | 0.954 | 0.908 | **0.893** | 0.786 | 0.841 | 0.702 |
| Class-weighted CE (Swin-B) | 0.946 | 0.913 | 0.881 | **0.810** | 0.829 | 0.687 |
| Mixup + BPaCo + cRT | 0.945 | 0.906 | 0.864 | 0.786 | 0.822 | 0.611 |

### S9. Bootstrap confidence intervals for the seed ensembles

Table S9
Three-seed probability-ensemble external macro-F1 with 95% confidence intervals from image-level nonparametric bootstrap resampling (2,000 replicates, percentile method). The intervals treat images as independent (no patient identifiers were available) and exclude seed variability (main Methods).

| Backbone | Ensemble macro-F1 | 95% CI |
|---|---|---|
| Swin (proposed) | **0.893** | 0.874–0.911 |
| VGG16-BN | 0.841 | 0.820–0.860 |
| ResNet50 | 0.751 | 0.737–0.766 |
| EfficientNet-B0 | 0.848 | 0.830–0.865 |

Source: `analysis/tables/external_macrof1_bootstrap_ci.csv`. VGG16-BN and EfficientNet-B0 rank here in the reverse of their seed-mean order in main-text Table 3 (0.813 versus 0.796): the two aggregations differ (probability ensemble versus mean over seeds); each ensemble estimate lies inside the other's 95% interval, and the seed-mean gap (0.017) is smaller than either across-seed sd.

### S10. Per-class external F1 by backbone

Table S10
Per-class F1 on the external benchmark by backbone (three-seed means), reported with the per-class external support $n$, which main-text Table 3 does not tabulate.

| Class | $n$ | Swin (proposed) | VGG16-BN | ResNet50 | EfficientNet-B0 |
|---|---|---|---|---|---|
| BASO | 74 | 0.899 | 0.883 | 0.870 | 0.897 |
| BLAST | 572 | **0.793** | 0.676 | 0.686 | 0.654 |
| EOS | 181 | 0.945 | 0.940 | 0.757 | 0.857 |
| LYM | 2,433 | 0.943 | 0.925 | 0.882 | 0.906 |
| MONO | 741 | 0.870 | 0.831 | 0.690 | 0.785 |
| NEU | 4,018 | 0.990 | 0.986 | 0.966 | 0.981 |
| PROMY | 42 | **0.601** | 0.446 | 0.177 | 0.491 |
| macro-F1 | 8,061 | **0.863** | 0.813 | 0.718 | 0.796 |
| weighted-F1 | | 0.947 | 0.927 | 0.886 | 0.911 |
| balanced acc | | 0.887 | 0.828 | 0.842 | 0.862 |

Source: `analysis/tables/master_external_results.csv`.

### Supplementary Note 1. Leave-one-dataset-out recipe and full-label-space results

The leave-one-dataset-out (LODO) models use the proposed recipe of Table S5, with seeds 42/52/62 in place of 42/62/82, so Table 4 characterizes the same classifier recipe the main results report, evaluated at the classifier level and without test-time augmentation. Each fold trains on the training splits of the four remaining sources (each source's own per-class 80/10/10 split), selects its checkpoint on their validation splits, leaves their test splits unused, and is tested on every pooled cell of the held-out source (Tables S1 and S15). An earlier arm on the identical folds and seeds used class-weighted cross-entropy *plus* mixup/CutMix (Zhang et al., 2018; Yun et al., 2019; $\alpha$ = 0.4); it scored lower on four of the five held-out domains (mean present-class macro-F1 0.685 versus 0.722; HRLS Lymphoblast F1 0.000 on every seed, as in the proposed arm, Table S14) and is retained in the code repository under `analysis/evidence/lodo/`. Per-seed values are given in `analysis/tables/lodo_corrected_perseed.csv`, whose nine-class column uses the union convention defined below. An earlier AML-LMU split omitted its typical lymphocytes (see the code repository); the corrected splits used here restore them. For the BMCMHF row, whose present-class macro-F1 has the widest spread (0.606 ± 0.030), the three per-seed values are 0.639 (seed 42), 0.612 (seed 52), and 0.566 (seed 62) — spread across all three seeds rather than one outlying run.

Main-text Table 4 reports present-class macro-F1; averaging F1 over the fixed nine-class label space, including zero-support classes, gives 0.525 ± 0.007 (AML-LMU), 0.606 ± 0.010 (PBC), 0.412 ± 0.008 (Raabin-WBC),

0.593 ± 0.006 (HRLS) and 0.471 ± 0.023 (BMCMHF), across-dataset mean 0.522 — 0.20 below the present-class mean purely because of zero-support classes. The denominator convention matters on one fold: averaging instead over the union of the true and predicted labels leaves four folds unchanged but raises PBC to 0.631 ± 0.027 and the mean to 0.526, because on seed 52 the model never predicts Lymphoblast, one of the three classes absent from PBC, so the denominator falls to eight.

### S11. Calibration and post hoc probability thresholds

In this post hoc analysis (main Methods), per-class one-vs-rest thresholds $t^\star$ for the three-seed probability ensemble were selected to maximize class F1 on the internal held-out test split ($n$ = 12,803) and applied unchanged to the external set ("internal-tuned"). Each threshold defines a binary decision — an image is positive for class $c$ if its seven-class ensemble probability $p_c \geq t$, regardless of the other classes — searched over $t = 0.01, 0.02, \ldots, 0.99$; the F1 values are one-vs-rest F1 of that decision, and no multiclass label or macro-F1 is recomputed. For reference we also report the oracle threshold tuned *on the external set itself* — an upper bound not attainable in deployment.

Table S11
Internal-tuned versus oracle (external-set-tuned) thresholds for the two clinically important minority classes. The internal-tuned threshold gives a small gain for PROMY but not for BLAST; the oracle column shows that even a perfectly tuned threshold would add little, so thresholding is not the main lever. "Internal F1" is measured on the split used to tune $t^\star$; Ext., external.

| Class | $t^\star$ (internal) | Internal F1 | Ext. F1 (argmax) | Ext. F1 (internal $t^\star$) | $t$ (oracle) | Ext. F1 (oracle) |
|---|---|---|---|---|---|---|
| BLAST | 0.500 | 0.949 | 0.841 | 0.839 | 0.380 | 0.849 |
| PROMY | 0.480 | 0.962 | 0.702 | 0.719 | 0.720 | 0.744 |

Source: `analysis/tables/threshold_validation_locked.csv`; BLAST oracle F1 from the counts in `analysis/evidence/wbcbench_confusion/threshold_sweep.csv.gz`.

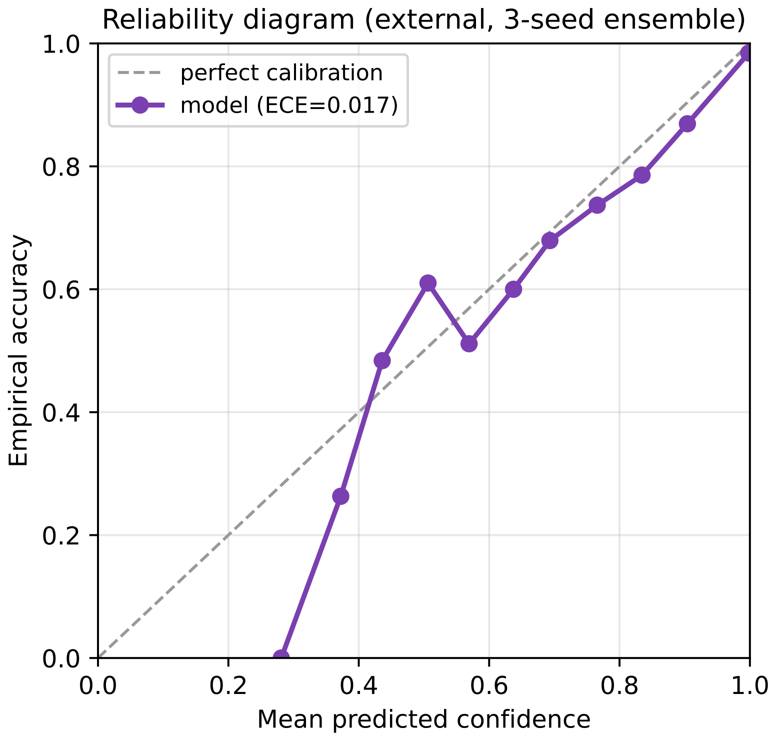


Fig. S2. Reliability diagram of the proposed model (three-seed ensemble) on the external WBCBench2026 Phase-1 set. Points show empirical accuracy versus mean predicted confidence per confidence bin (15 equal-width bins); the dashed line is perfect calibration. Expected calibration error was 0.017 and the multiclass Brier score 0.074 ($n$ = 8,061).

### S12. Are the detector's failures informative? A selective-prediction analysis

On the partly degraded Phase-2 split the detector returns no box for 348 labeled images (6.7%). The main text counts these as errors. This section asks whether that is too harsh: if the missed cells were simply unusable, discarding them would be a legitimate quality filter rather than a failure.

**Design.** Both groups were passed to the classifier *identically* (uncropped image → evaluation transform → three-seed Swin ensemble, no cropping), so the only difference in their treatment is whether the detector found a cell. The control group was matched on true class, which is essential: the missed set is 63% neutrophils, the easiest class, so an unmatched control would understate how much harder the missed cells are.

Table S12
Classification accuracy on the cells the detector missed, versus a class-matched control of cells it found, under identical input treatment. The missed cells are harder yet remain largely classifiable (interpretation below).

| Group | *n* | Accuracy | BLAST | EOS | LYM | MONO | NEU | PROMY |
|---|---|---|---|---|---|---|---|---|
| Missed by detector | 348 | **0.727** | 0.429 | 0.500 | 0.566 | **0.100** | 0.895 | 0.000 |
| Detected (class-matched) | 348 | **0.862** | 0.714 | 0.250 | 0.723 | 0.633 | 0.977 | 0.000 |
| Difference | | −0.135 | | | | | | |

Source: `analysis/tables/detector_abstention_quality_test.csv`; script `eval/detector_abstention_quality_test.py`. Per-class values are accuracy, over per-class group sizes (missed group: NEU 219, LYM 83, MONO 30, EOS 8, BLAST 7, PROMY 1; the class-matched control has the same sizes). The control was drawn once, and no significance test is reported.

**Interpretation.** Cells the detector misses are 13.5 accuracy points harder than class-matched cells it finds, and the steepest per-class drop is in monocytes (0.100 versus 0.633). The detector is therefore acting as an *implicit quality signal*: treating a no-detection as an abstention is a meaningful selective-prediction mechanism rather than an arbitrary exclusion.

We nevertheless report full coverage as the primary operating point: the classifier would still have labeled 73% of the abstained cells correctly, so discarding them would inflate the metric, and in deployment an abstained cell is routed to a human. Main-text Table 2 reports both operating points.

### S13. Two-hematologist review of hardest-class disagreements

Two hematologists at different hospitals independently reviewed the same 205 crops; the selection, review design and verdict categories are given in the main-text Results. Both are co-authors; neither had taken part in labeling the benchmark or read the manuscript before rating. The review form showed each crop (the detector's highest-confidence box at threshold 0.70, without margin) with the benchmark label and the model prediction, so the review was not blinded to either. The verdicts were not reconciled.

Table S13
Two-hematologist review of 205 hardest-class disagreements. Verdicts: *model right* (R), the benchmark label is wrong and the model is morphologically correct; *model wrong* (W), the benchmark label is right; *ambiguous* (A), difficult to distinguish or indeterminable. Disagreement types: Blast_FN, benchmark Blast group and model another class; Blast_FP, benchmark another (mature) class and model Blast group; Promy_FN and Promy_FP, likewise for Promyelocyte. The types partition the cells, with Blast group taking precedence, so Promy_FP here (15) excludes the four cells whose benchmark label is Blast group and whose model prediction is Promyelocyte (counted under Blast_FN); the main text's 19 Promyelocyte false alarms include them. (a) Verdicts of each hematologist by type. (b) Cross-tabulation of the two hematologists' verdicts; the diagonal holds the 73 shared verdicts (35.6%). (c) Paired outcomes by type; R–W counts cells rated R by one hematologist and W by the other, and likewise R–A and W–A.

*(a) Verdicts of each hematologist*

| | | First hematologist | | | Second hematologist | | |
|---|---|---|---|---|---|---|---|
| Type | *n* | R | W | A | R | W | A |
| Blast_FN | 93 | 57 | 30 | 6 | 12 | 72 | 9 |
| Blast_FP | 88 | 8 | 73 | 7 | 69 | 12 | 7 |
| Promy_FN | 9 | 7 | 2 | 0 | 0 | 9 | 0 |
| Promy_FP | 15 | 5 | 9 | 1 | 12 | 2 | 1 |
| Total | 205 | 77 (37.6%) | 114 (55.6%) | 14 (6.8%) | 93 (45.4%) | 95 (46.3%) | 17 (8.3%) |

*(b) Cross-tabulation*

| | Second hematologist | | | |
|---|---|---|---|---|
| First hematologist | R | W | A | Total |
| R | 21 | 50 | 6 | 77 |
| W | 69 | 43 | 2 | 114 |
| A | 3 | 2 | 9 | 14 |
| Total | 93 | 95 | 17 | 205 |

*(c) Paired outcomes*

| Type | *n* | R–R | W–W | A–A | R–W | R–A | W–A |
|---|---|---|---|---|---|---|---|
| Blast_FN | 93 | 11 | 29 | 4 | 42 | 5 | 2 |
| Blast_FP | 88 | 7 | 11 | 5 | 61 | 2 | 2 |
| Promy_FN | 9 | 0 | 2 | 0 | 7 | 0 | 0 |
| Promy_FP | 15 | 3 | 1 | 0 | 9 | 2 | 0 |
| Total | 205 | 21 (10.2%) | 43 (21.0%) | 9 (4.4%) | 119 (58.0%) | 9 (4.4%) | 4 (2.0%) |

Source: the two hematologists' per-cell verdicts, which are tied to WBCBench2026 images and labels and are not distributed.

**Reading.** The shared verdicts (diagonal of Table S13b) and the discordant pairs are reported in the main-text Results. Several of the 9 crops that both rated ambiguous showed only part of a cell, and a crop without margin gives little red-cell context for judging cell size. Most of the 119 R–W pairs were Blast_FN or Blast_FP cells (Table S13c). Individually, the first hematologist rated the model right on 77 (37.6%) and the second on 93 (45.4%).

### S14. Per-class leave-one-dataset-out performance

Table S14 decomposes each row of main-text Table 4 into its constituent classes. The present-class macro-F1 reported there is the unweighted mean of the F1 column within each block, so a class that fails completely is diluted by the others: AML-LMU scores 0.590 overall while its Myeloblast F1 is 0.048, and HRLS scores 0.763 while its Lymphoblast F1 is 0.000.

The two failures are not the same kind of failure. Of the 3,268 AML-LMU Myeloblasts, 78.5% were predicted Lymphocyte across the three seeds — a blast called a mature cell, which is the clinically costly direction. Of the 2,557 HRLS Lymphoblasts, 95.2% were predicted Myeloblast: the model recognizes the cell as a blast and assigns the wrong lineage. Because the harmonized seven-class space used for the external benchmark merges

Lymphoblast, Myeloblast and Erythroblast into a single Blast group, that second failure would not be scored as an error there.

The first failure is not a support artifact. The AML-LMU fold's training split held 2,027 HRLS myeloblasts (the training share of the 2,534 in Table S15; Supplementary Note 1), whereas the reverse fold, trained on 2,614 AML-LMU myeloblasts (and with no Lymphoblast training cells), reached HRLS Myeloblast recall 0.763. Common classes show the same pattern: Raabin-WBC Monocyte recall was 0.285 with 6,629 monocytes from the other four sources in training, and PBC Lymphocyte recall 0.666 with 23,749 lymphocytes. Supplying a class from other sources therefore did not ensure that it was recognized in a new source.

Table S14
Per-class leave-one-dataset-out performance of the proposed recipe (three-seed means, seeds 42/52/62) in the internal nine-class label space, with the evaluation setting of main-text Table 4. Classes are ordered by recall within each held-out source. **Bold** marks the two classes with recall below 0.10; their error routing differs in kind and is given in the text above.

| Held-out | Class | $n$ | Recall | F1 |
|---|---|---|---|---|
| AML-LMU | **Myeloblast** | 3,268 | **0.025** | **0.048** |
| | Basophil | 79 | 0.274 | 0.376 |
| | Monocyte | 1,779 | 0.765 | 0.779 |
| | Promyelocyte | 70 | 0.857 | 0.467 |
| | Erythroblast | 78 | 0.927 | 0.416 |
| | Lymphocyte | 3,947 | 0.947 | 0.710 |
| | Eosinophil | 424 | 0.952 | 0.947 |
| | Neutrophil | 8,483 | 0.994 | 0.982 |
| PBC | Lymphocyte | 1,214 | 0.666 | 0.724 |
| | Erythroblast | 1,547 | 0.781 | 0.876 |
| | Monocyte | 1,420 | 0.942 | 0.916 |
| | Basophil | 1,218 | 0.968 | 0.976 |
| | Eosinophil | 3,117 | 0.983 | 0.987 |
| | Neutrophil | 3,329 | 0.987 | 0.975 |
| Raabin-WBC | Monocyte | 561 | 0.285 | 0.431 |
| | Eosinophil | 744 | 0.767 | 0.730 |
| | Lymphocyte | 2,420 | 0.778 | 0.855 |
| | Basophil | 212 | 0.818 | 0.764 |
| | Neutrophil | 6,217 | 0.931 | 0.932 |
| HRLS | **Lymphoblast** | 2,557 | **0.000** | **0.000** |
| | Myeloblast | 2,534 | 0.763 | 0.548 |
| | Lymphocyte | 3,037 | 0.943 | 0.920 |
| | Monocyte | 2,039 | 0.965 | 0.893 |
| | Eosinophil | 1,017 | 0.987 | 0.993 |
| | Basophil | 1,023 | 0.999 | 0.993 |
| | Neutrophil | 3,198 | 0.999 | 0.993 |
| BMCMHF | Erythroblast | 13,470 | 0.272 | 0.421 |
| | Promyelocyte | 5,918 | 0.469 | 0.626 |
| | Basophil | 355 | 0.550 | 0.353 |
| | Monocyte | 3,049 | 0.671 | 0.379 |
| | Neutrophil | 26,536 | 0.825 | 0.864 |
| | Eosinophil | 3,484 | 0.832 | 0.838 |
| | Lymphocyte | 20,284 | 0.880 | 0.758 |

### S15. Per-dataset class composition of the leave-one-dataset-out pools

Table S15 decomposes each source dataset's harmonized single-cell pool by internal nine-class label. Each column is one dataset's complete pool, which is exactly the held-out set of that dataset's leave-one-dataset-out fold (Table S14); the support available to a fold for a class is therefore the Pooled column minus the held-out column, of which 80% was used for training and 10% for checkpoint selection (Supplementary Note 1). Three class–fold combinations lose 89–100% of their support this way: holding out HRLS leaves 0 of 2,557 Lymphoblast cells (the zero-shot fold of the main text), and holding out BMCMHF leaves 70 of 5,988 Promyelocyte and 1,625 of 15,095 Erythroblast cells — the two lowest-recall classes of that fold (0.469 and 0.272; Table S14) — so part of the BMCMHF fold's gap reflects the loss of class coverage that BMCMHF dominates (it holds 98.8% and 89.2% of the pooled Promyelocyte and Erythroblast cells) rather than bone-marrow morphology alone. Myeloblast, by contrast, splits 3,268/2,534 between AML-LMU and HRLS, so the AML-LMU fold retains 43.7% of its Myeloblast support; Section S14 shows that its Myeloblast failure is not a support artifact. Because each fold changes both the source and the class support, the folds do not rank the sources' marginal contributions to a common target.

Table S15
Per-dataset composition of the five harmonized single-cell pools, in the internal nine-class label space. Counts are cells surviving detector filtering at a confidence of 0.30 (0.70 for BMCMHF) and label harmonization (Tables S1 and S3). The merged internal corpus (127,938 cells; Table S2) is reconciled with these pools in Section S1. A zero means the class is absent from that source's pool after source-specific filtering and label mapping, not necessarily from the original dataset (HRLS Normoblast images, for example, were not mapped to Erythroblast; Section S3); the four classes below the rule are absent from at least one pool.

| | Held-out dataset | | | | | |
|---|---|---|---|---|---|---|
| Class | AML-LMU | PBC | Raabin-WBC | HRLS | BMCMHF | Pooled |
| Neutrophil | 8,483 | 3,329 | 6,217 | 3,198 | 26,536 | 47,763 |
| Lymphocyte | 3,947 | 1,214 | 2,420 | 3,037 | 20,284 | 30,902 |
| Monocyte | 1,779 | 1,420 | 561 | 2,039 | 3,049 | 8,848 |
| Eosinophil | 424 | 3,117 | 744 | 1,017 | 3,484 | 8,786 |
| Basophil | 79 | 1,218 | 212 | 1,023 | 355 | 2,887 |
| Erythroblast | 78 | 1,547 | 0 | 0 | 13,470 | 15,095 |
| Promyelocyte | 70 | 0 | 0 | 0 | 5,918 | 5,988 |
| Myeloblast | 3,268 | 0 | 0 | 2,534 | 0 | 5,802 |
| Lymphoblast | 0 | 0 | 0 | 2,557 | 0 | 2,557 |
| Total | 18,128 | 11,845 | 10,154 | 15,405 | 73,096 | 128,628 |

### S16. DinoBloom-B under leave-one-dataset-out

Table S16 reports DinoBloom-B fine-tuned on the identical leave-one-dataset-out folds, seeds, and present-class protocol as Table S14, in two configurations: the proposed recipe's hyperparameters (Table S5) applied to the DinoBloom-B backbone-specific model configuration (Section S7), and a BPaCo variant that combines the BPaCo term and augmentation of Section S7 with the proposed model's class-weighted cross-entropy and AdamW schedule. It exceeds the proposed model on every fold under both configurations (fold-equal grand means 0.776 and 0.759 versus 0.722; cell-weighted 0.715 and 0.692 versus 0.661). On three folds the comparison is confounded in the foundation model's favor: DinoBloom's self-supervised corpus (Koch et al., 2024) includes AML-LMU, Raabin-WBC and 80% of BMCMHF. PBC, which DinoBloom held out as its own external test set, and HRLS are not in that corpus; on these two folds DinoBloom-B still scored higher, by 0.009 and 0.018 with the proposed recipe and by 0.022 and 0.034 with the BPaCo variant. The pattern is consistent with breadth of exposure driving cross-domain robustness — for a self-supervised corpus as for the supervised multi-source pool — rather than proof of it. WBCBench2026, released after that corpus was assembled, is unseen by both models in training by provenance; there the proposed model scores higher on every matched seed (Section S7).

Table S16
DinoBloom-B under leave-one-dataset-out: same folds, seeds (42/52/62), classifier-level scope and present-class macro-F1 protocol as Table S14. Mean ± population sd over seeds. The two summary rows weight folds equally and by held-out cell count. Source: `analysis/evidence/lodo_foundation/`.

| | | | DinoBloom-B | |
|---|---|---|---|---|
| Held-out dataset | $n$ | Swin-B (proposed) | proposed recipe | BPaCo variant |
| AML-LMU | 18,128 | 0.590 ± 0.008 | 0.677 ± 0.004 | 0.647 ± 0.018 |
| PBC | 11,845 | 0.909 ± 0.015 | 0.918 ± 0.025 | 0.931 ± 0.011 |
| Raabin-WBC | 10,154 | 0.742 ± 0.015 | 0.844 ± 0.013 | 0.792 ± 0.034 |
| HRLS | 15,405 | 0.763 ± 0.008 | 0.781 ± 0.011 | 0.797 ± 0.000 |
| BMCMHF | 73,096 | 0.606 ± 0.030 | 0.660 ± 0.009 | 0.628 ± 0.017 |
| Mean, folds equal | | 0.722 | 0.776 | 0.759 |
| Mean, cell-weighted | | 0.661 | 0.715 | 0.692 |

### S17. Analysis specification and provenance

Table S17 lists, for each main classification analysis, the input route, the sample size, the evaluator and decision rule, the seed aggregation and the uncertainty estimate. It was compiled after the analyses from the code and output records. The three seed-aggregation rules are defined in the main Methods.

**Unified re-evaluation.** The Phase-1 route and Phase-2 results were first computed by three evaluation paths that differed in resize factor (1.1× or 1.14×), test-time augmentation and the order of the seven-class mapping. Every Phase-1 route and the Phase-2 split were re-scored post hoc with one evaluator — the common evaluator behind main-text Table 3. It reproduces the pre-cropped route and the uncropped-image route image for image; it changed 139–175 predictions in each of the runs first scored with the pipeline's own evaluator (route B and route E, seed 62; Phase-2, all seeds) and at most 51 in the others. The reported route and Phase-2 values are the unified ones. Macro-F1 seed means changed by −0.0025 for route E (0.8842 → 0.8817), by +0.002 for both Phase-2 views and by less than 0.001 elsewhere; route E Promyelocyte F1 changed most among the other reported values (0.698 → 0.680), and Phase-2 balanced accuracy rose from 0.786 to 0.795. Two analyses keep their own settings and were not re-derived: the resolution sweep of the main text (probability-summed mapping, except that its seed-62 run at 224 px is the common evaluator's; hence 0.863 ± 0.027 there, not 0.863 ± 0.028) and the missed-versus-detected comparison of Section S12.

**Development use of the external labels.** The proposed color-only policy was chosen from six augmentation policies, each replacing the default augmentation in the otherwise identical configuration of the class-weighted-CE baseline of Section S8: color-only (color jitter), stain-only (white-balance/gamma jitter), geometry-only (mild random-resized crop, flip, ±10° rotation), conservative (geometry-only plus milder color jitter), blur–noise (mild crop, flip, milder color jitter, light blur and Gaussian noise) and stain–blur–noise (blur–noise plus white-balance/gamma jitter). Their pre-cropped external macro-F1 (seed means, with flip averaging) ranged from 0.773 (stain–blur–noise) to 0.863 (color-only), against 0.856 for that baseline; flip averaging itself was adopted after comparing the same checkpoints without it (0.860 versus 0.863). The deployable route predates the cropper comparison of Section S6 and was not changed by it.

**Internal seven-class sensitivity.** Mapping the internal nine-class confusion matrices ($n$ = 12,803) to the seven evaluation classes gives internal macro-F1 of 0.952 (proposed Swin), 0.944 (VGG16-BN), 0.939 (EfficientNet-B0), 0.937 (ResNet50), 0.945 (Swin-B, BPaCo), 0.930 and 0.931 (ResNet50, BPaCo with AdamW and with SGD) and 0.953 (fine-tuned DinoBloom-B). The ordering among the four class-weighted-CE reference models, the position of DinoBloom-B and ResNet50's lower internal score under BPaCo with SGD than under class-weighted cross-entropy (0.931 versus 0.937) are the same as in the nine-class space.

Table S17
Analysis specification. Pre-cropped: highest-confidence detector box at 0.70, taxonomy-unmatched cells removed ($n$ = 8,061). Uncropped: all labeled uncropped images; the full pipeline runs the detector at 0.30. Common evaluator: square padding, resize to 256 px, center-crop to 224, horizontal-flip test-time augmentation. sd: population standard deviation over seeds.

| Quantity (where) | Input, $n$ | Evaluator and decision rule | Aggregation, uncertainty | Detection failures |
|---|---|---|---|---|
| Primary endpoint: full-pipeline macro-F1 (main Table 2, Phase-1 row) | Uncropped Phase-1, 8,080 | Common evaluator; per-seed argmax | Seed mean ± sd | Counted as errors (1 image) |
| Backbone comparison, external (main Table 3; S10) | Pre-cropped, 8,061 | Common evaluator; per-seed argmax | Seed mean ± sd | Not applicable |
| Internal macro-F1 (main Table 3) and blast sub-type F1 (S4) | Internal test, 12,803, nine classes | Each model's own evaluation preprocessing, no test-time augmentation | Seed mean (S4: ± sd) | — |
| Frozen probe (main Table 3) | Internal 12,803; pre-cropped 8,061 | Square padding, 256 px, crop 224 on every split; flip averaging externally | Single fit | Not applicable |
| Detection-and-cropping ablation and cropper routes (main Table 2; S6) | Uncropped Phase-1, 8,080 (route A: 8,061) | Common evaluator; per-seed argmax | Seed mean ± sd | Route B: counted; Cellpose routes classify every image |
| Phase-2 degradation (main Table 2) | Uncropped Phase-2, 5,196 (4,848 detected) | Common evaluator; per-seed argmax | Seed mean ± sd | Counted (348) or dropped (detected-only row) |
| Missed-versus-detected comparison (S12 table) | Phase-2 uncropped images, 348 + 348 | Resize 256, crop 224, no flip; three-seed probability sum, nine-class argmax mapped | Descriptive, single draw | Missed cells are the subject |
| Other ResNet50 and Swin-B configurations (main text; S7) | Pre-cropped, 8,061 | Common evaluator; per-seed argmax | Seed mean ± sd | Not applicable |
| Recipe-selection bootstrap (main text; S8) | Pre-cropped, 8,061 | Common evaluator; probability ensembles | Paired image bootstrap, 1,000 replicates, run during recipe selection | Not applicable |
| Ensemble macro-F1 intervals (S9; main text) | Pre-cropped, 8,061 | Common evaluator; probability ensemble | Image bootstrap, 2,000 replicates | Not applicable |
| Ensemble point estimates (S8), calibration, confusion matrices, error analysis | Pre-cropped, 8,061; internal matrix: internal test, 12,803 | Common evaluator; probability ensemble | No interval | Not applicable |
| Paired tests (S7) | Pre-cropped, 8,061 | Common evaluator; majority vote | McNemar; paired bootstrap, 5,000; TOST at 90% | Not applicable |
| Seed-level test, proposed vs. fine-tuned DinoBloom-B (S7) | Pre-cropped, 8,061 | Common evaluator; per-seed argmax | Paired $t$-test, three seed pairs | Not applicable |
| Leave-one-dataset-out (main Table 4; S14, S16) | Held-out source cells passing the detector filter at 0.30 (0.70 for BMCMHF) | Common evaluator without test-time augmentation; nine classes, present-class macro-F1 | Seed mean ± sd (42/52/62) | Not counted (conditional) |
| Threshold analysis (S11; post hoc) | Tuned on internal test, 12,803; applied to pre-cropped, 8,061 | Common evaluator; one-vs-rest thresholds | Probability ensemble; no interval | Not applicable |
| Resolution sweep (main text) | Pre-cropped, 8,061 | Flexible-resolution evaluator; mapping: see text above | Seed mean ± sd | Not applicable |
